# Collagen VI regulates motor circuit plasticity and motor performance by cannabinoid modulation

**DOI:** 10.1101/2021.09.03.458830

**Authors:** Daniel D. Lam, Rhîannan H. Williams, Ernesto Lujan, Koji Tanabe, Georg Huber, Nay Lui Saw, Juliane Merl-Pham, Aaro V. Salminen, David Lohse, Sally Spendiff, Melanie J. Plastini, Michael Zech, Hanns Lochmüller, Arie Geerlof, Stefanie M. Hauck, Mehrdad Shamloo, Marius Wernig, Juliane Winkelmann

**Affiliations:** Institute of Neurogenomics, Helmholtz Center Munich, German Research Center for Environmental Health, 85764 Neuherberg, Germany; Chair of Neurogenetics, Neurological Clinic and Polyclinic, Klinikum rechts der Isar, School of Medicine, Technical University of Munich, 81, 81675 Munich, Germany; Institute for Stem Cell Biology and Regenerative Medicine, Department of Pathology, Stanford University, Stanford, CA 94305, USA; Protein Expression and Purification Facility, Institute of Structural Biology, Helmholtz Center Munich, German Research Center for Environmental Health, 85764 Neuherberg, Germany; Department of Neurosurgery, Stanford University School of Medicine, Palo Alto, CA 94304, USA; Research Unit Protein Science, Helmholtz Center Munich, German Research Center for Environmental Health, 85764 Neuherberg, Germany; Children’s Hospital of Eastern Ontario Research Institute, Ottawa, ON K1H 5B2, Canada; Department of Neurology and Neurological Sciences, Stanford University School of Medicine, Palo Alto, CA 94304, USA; Division of Neurology, Department of Medicine, The Ottawa Hospital, Ottawa, ON K1H 8L6, Canada; Brain and Mind Research Institute, University of Ottawa, Ottawa, ON K1N 6N5, Canada; Munich Cluster for Systems Neurology (SyNergy), Munich, Germany; Institute of Human Genetics, Klinikum rechts der Isar, School of Medicine, Technical University of Munich, 81675 Munich, Germany

## Abstract

Collagen VI is a key component of muscle basement membranes, and genetic variants can cause monogenic muscular dystrophies. Conversely, human genetic studies recently implicated collagen VI in central nervous system function, with variants causing the movement disorder dystonia. To elucidate the neurophysiological role of collagen VI, we generated mice with a truncation of the dystonia-related collagen α3 (VI) (COL6A3) C-terminal domain (CTD). These *Col6a3*^CTT^ mice showed a recessive dystonia-like phenotype. We found that COL6A3 interacts with the cannabinoid receptor 1 (CB1R) complex in a CTD-dependent manner. *Col6a3*^CTT^ mice have impaired homeostasis of excitatory input to the basal pontine nuclei (BPN), a motor control hub with dense COL6A3 expression, consistent with deficient endocannabinoid signaling. Aberrant synaptic input in the BPN was normalized by a CB1R agonist, and motor performance in *Col6a3*^CTT^ mice was improved by CB1R agonist treatment. Our findings identify a readily therapeutically addressable synaptic mechanism for motor control.

**SIGNIFICANCE STATEMENT:** Dystonia is a movement disorder characterized by involuntary movements. We previously identified genetic variants affecting a specific domain of the COL6A3 protein as a cause of dystonia. Here, we we created mice lacking the affected domain and observed an analogous movement disorder. Using a protein interaction screen, we found that the affected COL6A3 domain mediates an interaction with the cannabinoid receptor CB1R. Concordantly, our COL6A3-deficient mice showed a deficit in synaptic plasticity linked to a deficit in cannabinoid signaling. Pharmacological cannabinoid augmentation rescued the motor impairment of the mice. Thus, cannabinoid augmentation could be a promising avenue for treating dystonia, and we have identified a possible molecular mechanism mediating this.

## INTRODUCTION

Collagen VI forms supramolecular microfibrillary networks in the extracellular matrix (Timpl and Chu, 1994). It is a major component of muscle basement membrane, where it supports muscle stiffness and elasticity structurally as well as by regulating cellular processes (Grumati et al., 2010; Irwin et al., 2003; Urciuolo et al., 2013). Variants in collagen VI-encoding genes can cause monogenic muscular dystrophies due to the integral role of collagen VI in muscle function (Bönnemann, 2011).

Collagen VI is assembled from three subunits, one of which is collagen α3 (VI) or COL6A3 (Timpl and Chu, 1994) (Fig. 1a). We recently identified bi-allelic point and indel mutations of the unique COL6A3 C-terminal domain (CTD) in early-onset dystonia (Zech et al., 2015), a movement disorder characterized by sustained or paroxysmal involuntary abnormal movements and/or postures (Albanese et al., 2013). This suggests a CTD-dependent function of COL6A3 in the central nervous system.

**Figure 1.**
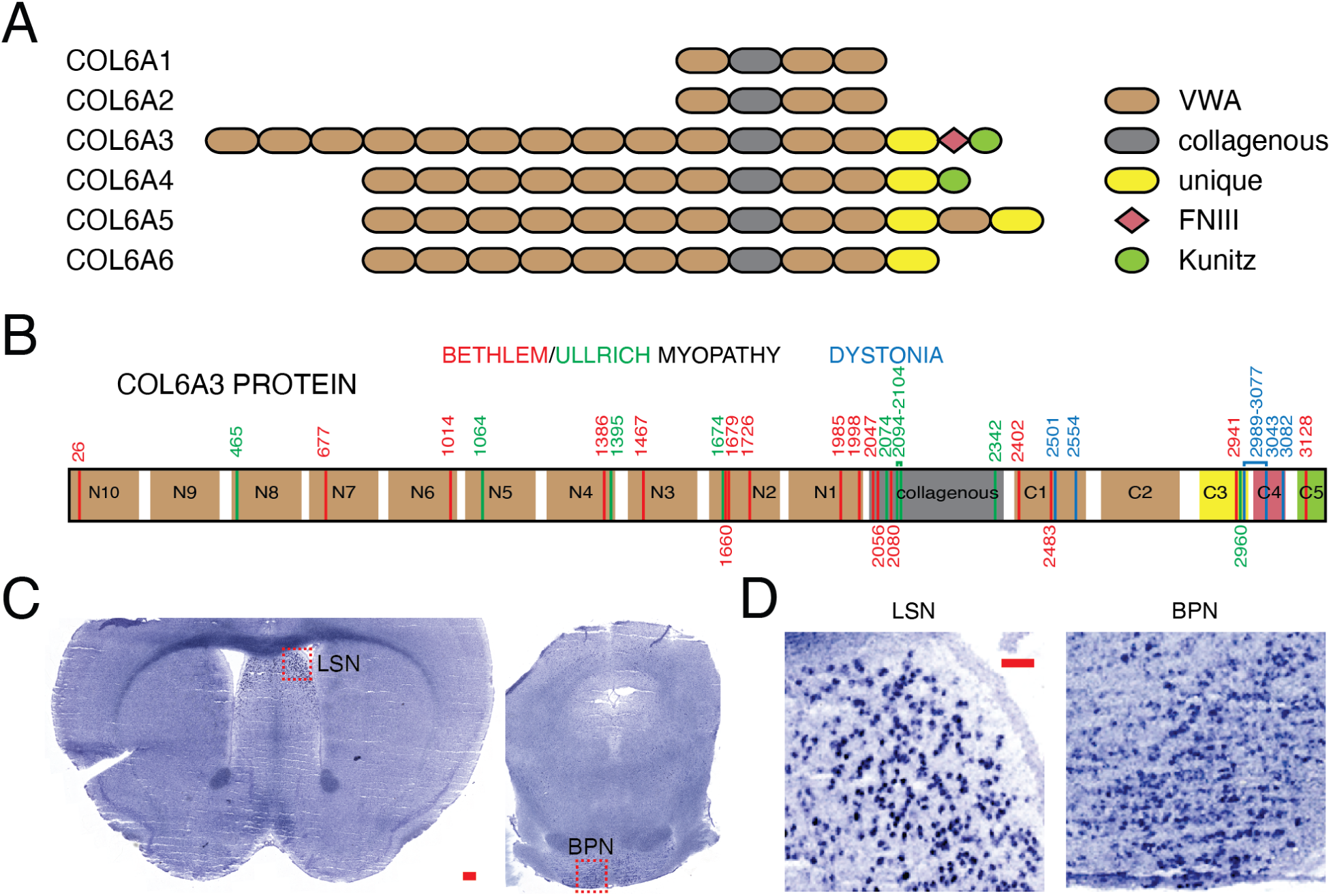
Domains and expression of COL6A3. (**A**) Domains of collagen VI subunits. VWA, von Willebrand Factor Type A; FNIII, fibronectin type III. (**B**) Domains of COL6A3, including globular N-terminal domain (N1-10), collagenous domain, and globular C-terminal domain (C1-5). Indicated are amino acid positions of disease-associated variants from UniProt and other reports (Briñas et al., 2010; Demir et al., 2002; Marakhonov et al., 2018; Shimomura et al., 2019; UniProt Consortium, 2019; Zhang et al., 2014). (**C**) Primary sites of adult mouse brain *Col6a3* expression (dark purple) by in situ hybridization in coronal sections. Highlighted are the lateral septal nucleus (LSN) and basal pontine nuclei (BPN), where expression is most concentrated. Scale bar, 200 µm. (**D**) Magnification of boxed areas in (C). Scale bar, 50 µm.

To unravel the mechanism linking the COL6A3 CTD with dystonia, we generated *Col6a3*^CTT^ mice with a truncation of this domain and observed a recessive dystonia-like phenotype. Probing the molecular mechanism, we identified a CTD-dependent interaction between COL6A3 and the cannabinoid receptor 1 (CB1R) complex. The endocannabinoid (eCB) system restrains synaptic input and maintains synaptic homeostasis via inhibitory retrograde signaling (Wilson and Nicoll, 2001). In *Col6a3*^CTT^ mice, this process was impaired in neurons of the basal pontine nuclei, a motor control interface between motor cortex and cerebellum with dense COL6A3 expression. This synaptic deficit, leading to inadequate synaptic downscaling, is a plausible mechanism for hallmarks of dystonia including the central hyperkinetic aspect, impaired surround inhibition, and task-specific manifestations (Calabresi et al., 2016; Quartarone and Pisani, 2011).

Despite these synaptic deficits, CB1R remained intact and could be targeted with an agonist *in vivo*, where this treatment significantly improved motor performance in *Col6a3*^CTT^ mice. Thus, cannabinoid augmentation represents a promising noninvasive therapeutic modality with robust neurophysiological evidence.

## RESULTS

### Domains and expression of COL6A3

Collagen α3 (VI), or COL6A3, consists of globular N and C terminal domains, which are divided into subdomains (N1-10 and C1-5) on the basis of structural motifs, and a collagenous, or triple helical, domain (Figure 1A). Uniquely among collagen VI subunits, COL6A3 contains a fibronectin type III C4 domain (Bonaldo and Colombatti, 1989), which is affected by the majority of dystonia-related variants (Zech et al., 2015) (Figure 1B). Indeed, each of the *COL6A3*-related dystonia families harboured at least one deleterious variant in the portion of this domain encoded by exon 41 (Zech et al., 2015) (Figure 1B). In contrast, Bethlem and Ullrich myopathy-related variants almost exclusively affect the N-terminal and collagenous domains (Figure 1B).

Given the paucity of information, we asked whether *Col6a3* is expressed in the brain. *In situ* hybridization experiments in mouse brain revealed that *Col6a3* expression is highly restricted and is only found in two notable clusters, in the lateral septal nucleus (LSN) and the basal pontine nuclei (BPN) (Figures 1C and 1D).

### COL6A3 attenuation causes recessive dyskinesia

To delineate the function of the COL6A3 C-terminal domain (CTD), which is selectively affected in dystonia patients, we generated COL6A3 C-terminal truncation (*Col6a3*^CTT^) mice. To that end, we inserted a premature stop codon into exon 36, encoding the C1 domain (Figure S1A). This mutation, verified at the RNA level (Figure S1B), is expected to result in a truncated COL6A3 protein lacking the majority of the CTD. Both heterozygous and homozygous *Col6a3*^CTT^ mice were born in Mendelian ratios, were viable and fertile, and did not show any gross morphological abnormalities.

COL6A3 expression was reduced in *Col6a3*^CTT/CTT^ mice, presumably due to nonsense-mediated decay (Figure S1C-H). Based on RNA-seq analysis, wildtype mice express an mRNA encoding full-length COL6A3 in the BPN, where expression is ∼25% of the level of tibialis anterior muscle (Figure S1D and Figure S2). COL6A3 protein levels generally reflected mRNA levels (Figure S1H). In *Col6a3*^CTT/CTT^ mice, exon usage and splicing were indistinguishable from wildtype (Figure S2). At age 2 months, *Col6a3*^CTT/CTT^ mice show no overt (but possibly incipient) muscle pathology, while older mice develop a degenerative myopathy of modest severity (Figure S3 and Figure S4). Specifically, 2 month old mice show no significant change in the proportion of internalised nuclei, a sign of damaged myofibres (Folker and Baylies, 2013). In contrast, 13 month old mice had more internalised nuclei and greater variability in myofibre size and shape, indicative of ongoing degeneration (Figure S3).

Upon behavioural characterization, *Col6a3*^CTT^ mice showed a marked recessive dyskinetic phenotype, characterised in the beam walk test by increased slips (wildtype 1.9 ± 0.4 slips; *Col6a3*^CTT/CTT^ 4.0 ± 0.6 slips; Welch *t*-test: p = 0.015, t = −2.7, df = 17.3; Figures 2A and 2B), profound hindlimb incoordination (wildtype 0.03 ± 0.03 trials; *Col6a3*^CTT/CTT^ 0.96 ± 0.25 trials; Welch *t*-test: p=0.001, t = −3.6, df = 24.6; Figure 2C and Movie S1), and slower traversal speed (wildtype 9.0 ± 0.4 sec; *Col6a3*^CTT/CTT^ 11.4 ± 0.8 sec; Welch *t*-test: p = 0.008, t = −2.8, df = 38.3; Figure 2D). The recessive nature of the phenotype accords with the recessive inheritance pattern of dystonia caused by COL6A3 CTD variants in humans (Zech et al., 2015). *Col6a3*^CTT/CTT^ (homozygous) mice produced less grip strength than wildtypes (wildtype 1.4 ± 0.1 N; *Col6a3*^CTT/CTT^ 1.1 ± 0.1 N; Welch *t*-test: p = 0.001, t = 3.5, df = 37.9; Figure 4E). *Col6a3*^CTT^ mice also showed a significant recessive impairment on the rotarod (Figure 4F), while locomotor performance (distance and speed) were not impaired (Figures 4G and 4H). The behavioural phenotype was consistent across different ages, laboratories, and assessors. One cohort of females aged 7-8 wk was characterized at the Stanford Behavioral and Functional Neuroscience Laboratory, where total slips on the 12 mm beam were 1.9 ± 0.4 slips for wildtypes and 4.0 ± 0.6 for *Col6a3*^CTT/CTT^; p=0.0015; see above). Another cohort of males and females aged 12-28 wk was characterized at the Helmholtz Centre Munich, where the corresponding values were 1.0 ± 0.2 slips for wildtypes and 2.2 ± 0.5 for *Col6a3*^CTT/CTT^; Welch *t*-test: p=0.03, t = −2.2, df = 40.1).

**Figure 2.**
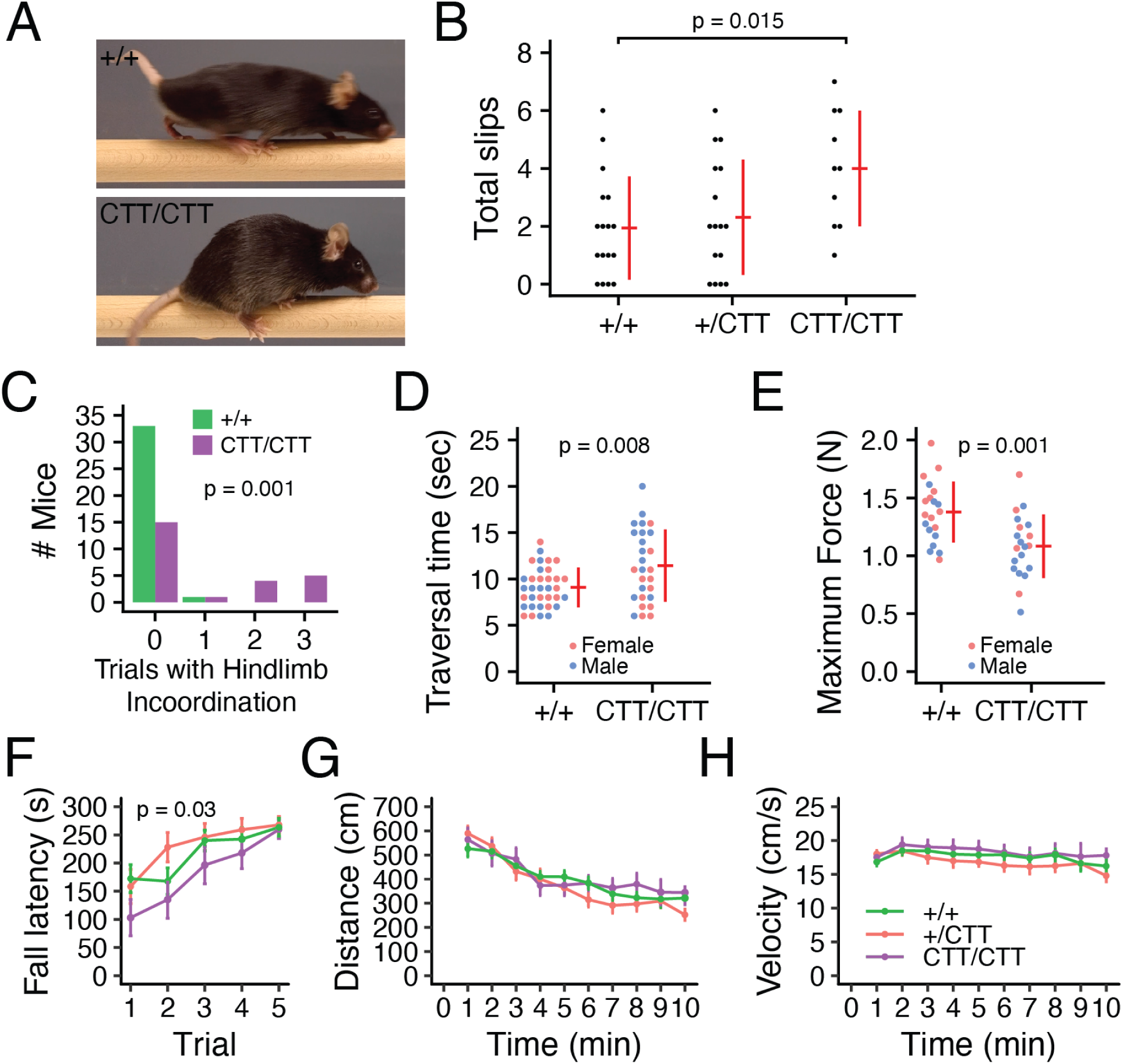
COL6A3 attenuation causes recessive dyskinesia. (**A**) Representative images of *Col6a3*^+/+^ and *Col6a3*^CTT/CTT^ mice during beam walk, showing the frame immediately before right forepaw liftoff. (**B**) Total hindlimb slips across three beam walk trials (12 mm rectangular beam). Female mice aged 7-8 wk, n=10-17 per genotype. (**C**) Occurrence of hindlimb incoordination across three beam walk trials (12 mm rectangular beam). Male and female mice aged 12-28 wk, n=25-34 per genotype. (**D**) Best traversal time from 3 beam walk trials (12 mm rectangular beam). Male and female mice aged 12-28 wk, n=27-34 per genotype. (**E**) Maximum four limb force in grip strength test. Male and female mice aged 23-29 wk, n=20 per genotype. (**F-H**) Rotarod fall latency on 5 consecutive trials (F); distance traveled (G) and velocity (H) per minute in activity chamber. Female mice aged 7-8 wk, n=10-17 per genotype.

All error bars are standard deviation and centre values are group means. *P* values derived from t-test, except for (F), one-way ANOVA between wildtypes and *Col6a3*^CTT/CTT^. See also Figures 2-1, 2-2, 2-3, 2-4, and Video 2-1.

### COL6A3 interacts with cannabinoid receptor 1 (CB1R)

To identify binding partners of the COL6A3 C4 domain, the domain most impacted by dystonia variants, we performed an unbiased protein interaction screen. We first synthesized full-length wildtype murine COL6A3 (COL6A3^WT^) and COL6A3 with the C4 domain deleted (COL6A3^ΔC4^). The proteins were synthesized using baculovirus in insect cells (Figure S5). As proline hydroxylation is critical for posttranslational folding and normal conformation of collagen molecules (Chopra and Ananthanarayanan, 1982; Vitagliano et al., 2001), we concomitantly expressed proline hydroxylase. The proteins were then coupled to beads and incubated with mouse brain lysates followed by mass spectrometry of the bead eluate (Figure 3A).

**Figure 3.**
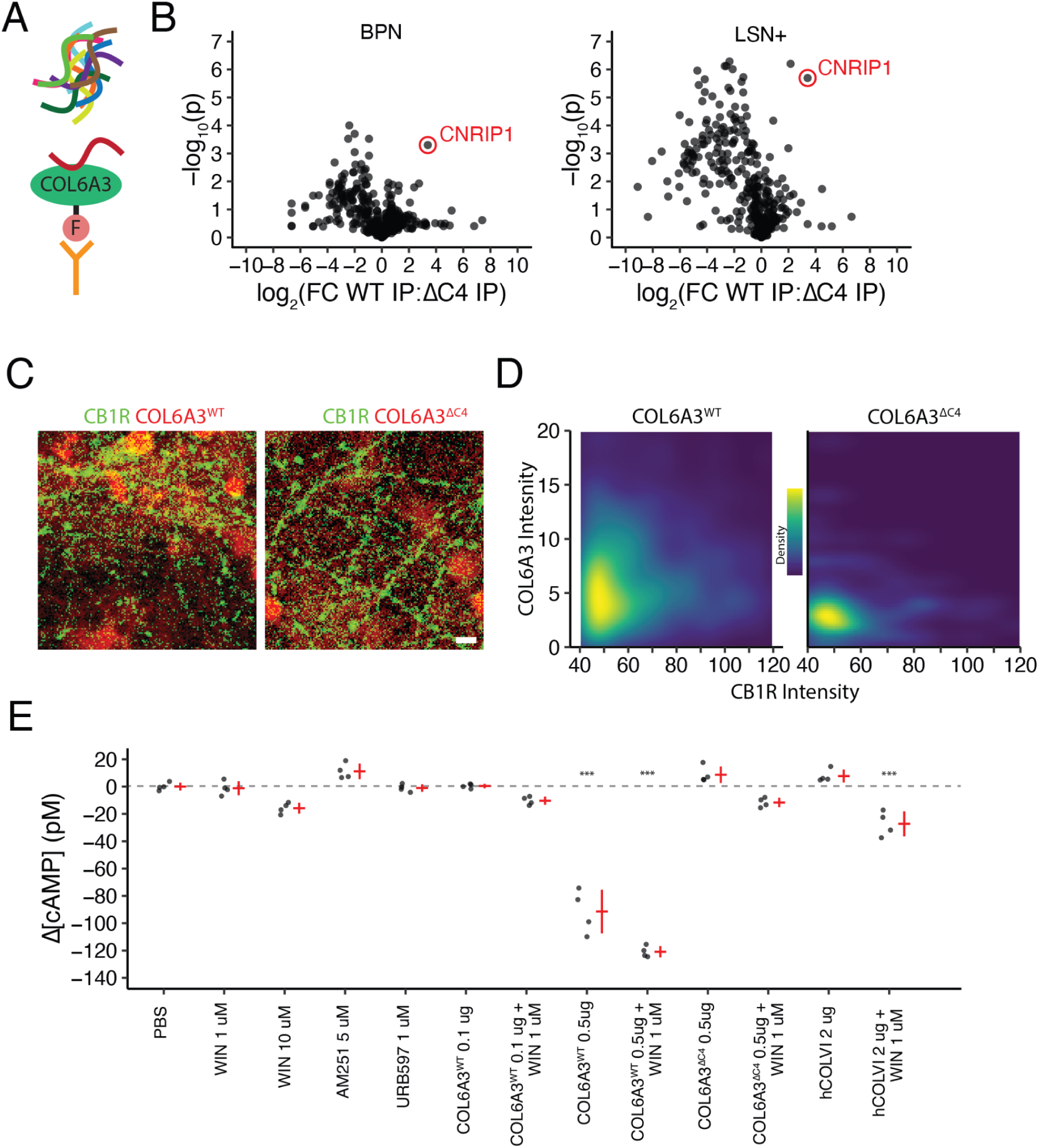
COL6A3 interacts with cannabinoid receptor 1. (**A**) Schematic of immunoprecipitation (IP) experiment. FLAG-tagged COL6A3 was conjugated to beads with anti-FLAG antibody, incubated with mouse brain lysates, and eluate was analysed with mass spectrometry. (**B**) Results of mass spectrometry analysis of IP from BPN and LSN lysates. Each point is a protein. The abscissa is the base 2 logarithm of the ratio of abundance of each protein in the COL6A3^WT^ to COL6A3^ΔC4^ IP, and the ordinate is the negative base 10 logarithm of the p value comparing the two IPs (two-sided *t*-test, not corrected for multiple comparisons). Proteins towards the top right bind selectively to COL6A3^WT^ compared to COL6A3^ΔC4^. (**C**) Representative images from wash-and-bind assay on primary mouse cortical neurons showing colocalization (yellow) of COL6A3^WT^ (red) with CB1R (green). Scale bar, 10 µm. (**D**) Pixel-wise density plots of wash-and-bind results, comparing CB1R fluorescence with retention of COL6A3^WT^ or COL6A3^ΔC4^ (detected by anti-His immunofluorescence). Results are from 3 technical replicates per condition. (**E**) cAMP assay in mouse primary neurons. COL6A3^WT^, COL6A3^ΔC4^, or human collagen VI preparation were added to cells alone or in combination with CB1R agonist WIN 55,212-2 (WIN). All error bars are standard deviation and centre values are group means. *P* values derived from ANOVA followed by post-hoc Tukey HSD test comparing to PBS, *** p < 0.001. See also Figure 3-1.

Using both BPN and LSN lysates from microdissected mouse brains, this approach identified the cannabinoid receptor interacting protein CNRIP1 as preferentially binding to the C4 domain of COL6A3 (Figure 3B). As membrane proteins are difficult to access with detergent solubilization, we surmised that this association occurs between the extracellular COL6A3, the membrane-integral CB1R, and the intracellular CNRIP1.

To independently confirm the COL6A3-CB1R interaction, we performed wash-and-bind assays in primary cortical neurons, showing that COL6A3 binding was positively correlated with extent of CB1R expression (Figure 3C and 3D). We performed Pearson’s correlations between COL6A3 (WT or ΔC4) and CB1R fluorescence for each pixel and obtained r^2^ values of 0.05 for COL6A3^WT^ and −0.006 for COL6A3^ΔC4^. Comparison of these correlations by Fisher’s z transformation revealed a significant effect of C4 domain deletion on the correlation of CB1R fluorescence with COL6A3 retention, indicating that the retention of COL6A3^ΔC4^ by CB1R was significantly reduced compared to COL6A3^WT^ (p = 0.02).

Finally, we showed that COL6A3^WT^, but not COL6A3^ΔC4^, synergized with CB1R agonist to reduce cAMP levels (Figure 3E). Specifically, the effect of 0.5 µg COL6A3^WT^ + WIN was greater than for COL6A3^WT^ alone (ANOVA with post-hoc Tukey HSD, p = 0.0001). The effect of a human collagen VI preperation was also greater with WIN than without (ANOVA with post-hoc Tukey HSD, p = 3 x 10^-6^).

### COL6A3 is produced by a distinct pontine cell type

To determine COL6A3-expressing cell types, we harnessed a recently published single cell RNA sequencing catalogue of mouse nervous system cells (Zeisel et al., 2018). The central nervous system cell clusters with the highest *Col6a3* trinarization score, a probabilistic estimate of gene expression in cell types, were excitatory brainstem neurons mapping to the BPN and inhibitory neurons of the septal nucleus (Figure 4A). These results independently confirm our *in situ* hybridization findings (Figure 1C and 1D). Within the 97 hindbrain neurons expressing *Col6a3*, expression of *Col6a3* was positively correlated with the excitatory neuron marker *Slc17a7* (Vglut1) and negatively correlated with the inhibitory neuron markers *Gad1* and *Slc32a1* (Vgat) (Figure 4B), indicating that *Col6a3* expression is specific to excitatory neurons over inhibitory neurons. *In situ* hybridization confirmed that BPN *Col6a3*-expressing cells predominantly co-express the glutamatergic neuron marker *Slc17a7* (89 ± 3%, n=3; Figure 4C). BPN *Col6a3*-expressing cells also predominantly co-express *Dagla* (84 ± 4%, n=3; Figure 4C), which encodes diacylglycerol lipase ɑ, the catalytic enzyme for biosynthesis of the endocannabinoid 2-arachidonoylglycerol (2-AG). With the caveat that RNA expression does not confirm protein expression, we conclude that BPN glutamatergic neurons can synthesize both collagen VI and 2-AG.

**Figure 4.**
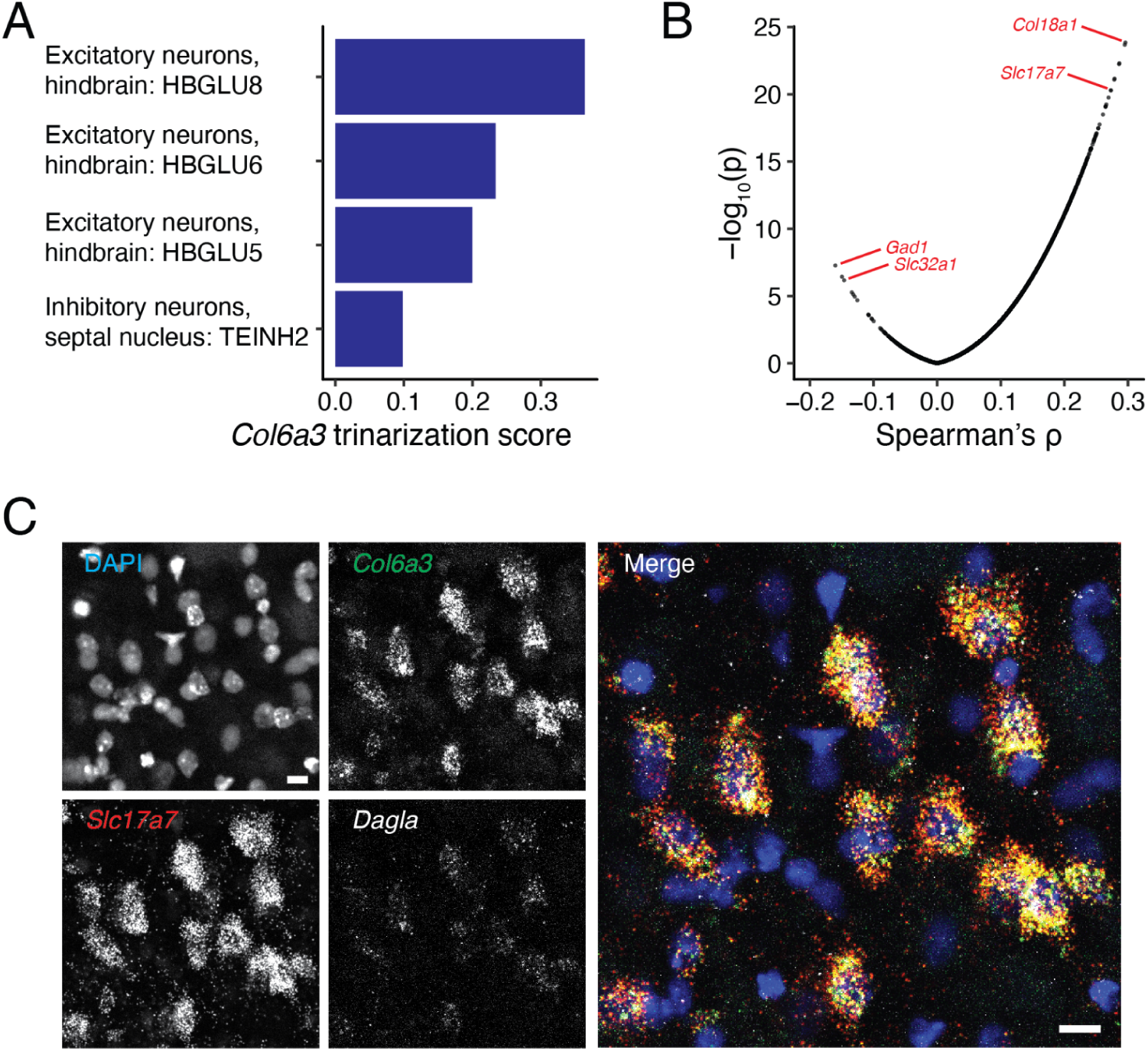
COL6A3 is produced by a distinct pontine cell type. (**A**) Ranking of central nervous system cell clusters by *Col6a3* trinarization score (a probabilistic cluster gene expression estimate), using data from Zeisel et al., 2018. (**B**) Transcriptome-wide analysis of coexpression with *Col6a3* reveals positive correlation of the glutamatergic neuron marker *Slc17a7* (*Vglut1*) and negative correlation of the inhibitory neuron markers *Gad1* and *Slc32a1* (*Vgat*), indicating that *Col6a3*-expressing cells are glutamatergic neurons. (**C**) Triple *in situ* hybridization of adult mouse BPN showing predominant coexpression of *Slc17a7* and the endocannabinoid synthesis gene *Dagla* in *Col6a3*-expressing cells. Scale bars, 20 µm.

### COL6A3 truncation impairs plasticity of pontine afferents

Since COL6A3 interacts with CB1R, which mediates synaptic downscaling *via* retrograde inhibition (Wilson and Nicoll, 2001), we hypothesized that COL6A3 truncation may affect synaptic plasticity. Specifically, we hypothesized that COL6A3 truncation would perturb long-term synaptic depression mediated by CB1R (Chevaleyre and Castillo, 2003; Kreitzer and Malenka, 2007; Robbe et al., 2002). We reasoned that collagen VI would, analogous to collagen IV, affect afferent axons at the site of collagen secretion (Xiao and Baier, 2007; Xiao et al., 2011). Further justifying the assumption that collagen VI is secreted locally, the *Col6a3*^CTT/CTT^ genotype had a much more significant effect on the BPN proteome than that of the cerebellum (Figure S6).

Thus, we performed *ex vivo* whole-cell patch-clamp electrophysiological recordings on BPN neurons of acute brain slices, around which collagen VI is expected to be secreted (Figure 5A and 5B). The BPN consists of exclusively glutamatergic neurons (Figure 5C), primarily conveying descending signals from the cortex to the cerebellum (Brodal and Bjaalie, 1997; Kelly and Strick, 2003; Suzuki et al., 2012). We found that the intrinsic passive membrane properties, namely resting membrane potential (RMP), cell capacitance and input resistance (R_in_; Table S1), and both glutamatergic and GABAergic tone onto these cells were similar in wildtype and *Col6a3*^CTT/CTT^ mice (Figure 5D-M). A subset of *Col6a3*^CTT/CTT^ neurons did have higher spontaneous excitatory postsynaptic current (sEPSC) activity, yet as a population it did not reach statistical significance (Figure 5I, 5J, and 5L). Action potential properties were also similar between genotypes (Table S1).

**Figure 5.**
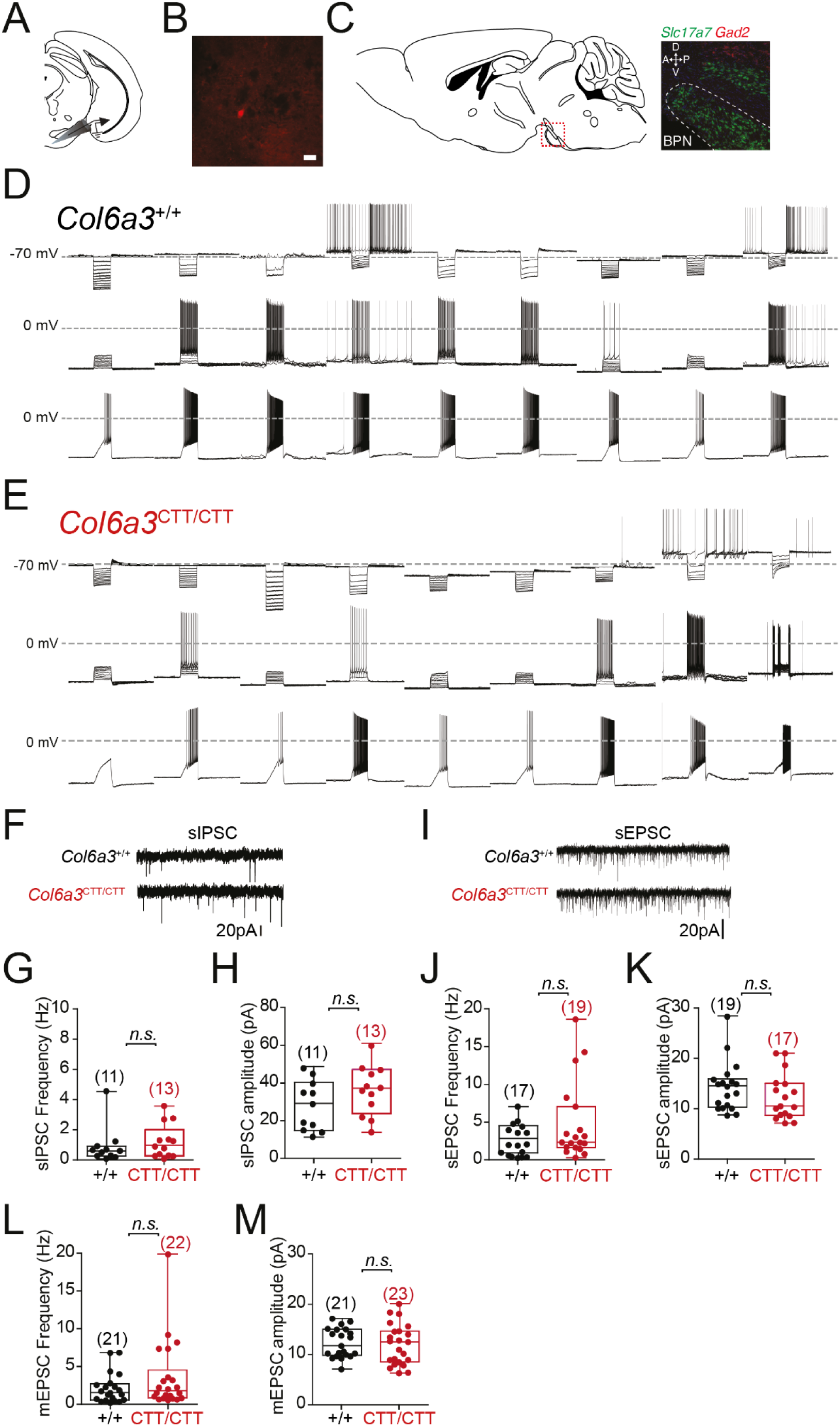
Intrinsic properties of BPN neurons. (**A**) Location of patch clamp recordings. Schematic of coronal brain section with targeted region indicated with electrode. (**B**) Example of patched BPN neuron. The recorded cell was filled with neurobiotin and immunolabeled with streptavidin-Alexa Fluor 594. Filled cell visible in red. Scale bar, 20 µm. (**C**) BPN cell type. Left, schematic sagittal section with red box indicating position of detail. Right, *in situ* hybridization showing *Slc17a7* (glutamatergic neuron marker) in green, *Gad2* (GABAergic neuron maker) in red, and DAPI (nuclei) in blue. All BPN neurons are glutamatergic. (**D,E**) Representative current-clamp recordings of different wildtype (D) and *Col6a3*^CTT/CTT^ (E) BPN neurons showing mixture of spontaneously firing and silent cells and their responses to current injection (top rows: −20 pA steps; second rows: +10 pA steps; bottom rows: +200 pA ramp). Each column is from the same cell. (**F**) Representative voltage-clamp recordings of wildtype and *Col6a3*^CTT/CTT^ BPN neurons showing spontaneous inhibitory postsynaptic currents (sIPSC). (**G,H**) Spontaneous inhibitory postsynaptic current frequency (G) and amplitude (H). (**I**) Representative voltage-clamp recordings of wildtype and *Col6a3*^CTT/CTT^ BPN neurons showing spontaneous excitatory postsynaptic currents (sEPSC). (**J,K**) Spontaneous excitatory postsynaptic current frequency (J) and amplitude (K). (**L,M**) Miniature excitatory postsynaptic current frequency (L) and amplitude (M). See also Table 5-1 and Figure 5-1.

The type I metabotropic glutamate receptor (mGluR) agonist (S)-3,5-dihydroxyphenylglycine (DHPG) drives eCB-mediated inhibition of presynaptic inputs (Chevaleyre and Castillo, 2003; Kushmerick et al., 2004; Robbe et al., 2002; Varma et al., 2001). We therefore evaluated this response in BPN neurons of wildtype and *Col6a3*^CTT/CTT^ mice. DHPG (100µM) applied in the presence of tetrodotoxin (TTX, 1 µM; to prevent action potential-mediated responses) caused a large significant inward current in both genotypes when compared to baseline values (WT: −36.68 ± 5.17 pA, n=16; and *Col6a3*^CTT/CTT^: −49.79 ± 11.22 pA, n=13). The current evoked in *Col6a3*^CTT/CTT^ cells failed to return to baseline values despite a prolonged washout period (>20min), suggesting altered mGluR kinetics (Figure 6A and 6A”). The current was largely and significantly attenuated by the CB1R agonist WIN 55,212-2 (WIN, 1µM;), (WT:+DHPG −47.43 ± 9.03 pA, and +WIN/DHPG −12.77 ± 5.67 pA, n=6, p = 0.007; and *Col6a3*^CTT/CTT^:+DHPG −29.57 ± 6.8 pA, and +WIN/DHPG −3.66 ± 2.34 pA, n=5, p = 0.005; Figre 6B, 6B’, 6C, and 6C’), presumably via inhibition of presynaptic glutamate release. The attenuation of DHPG current by WIN was similar between genotypes (wildtype, 75.7 ± 12.7% reduction, n=6; CTT/CTT, 92.5 ± 6.0% reduction, n=5; p=0.31; Figure 6C and 6C’).

**Figure 6.**
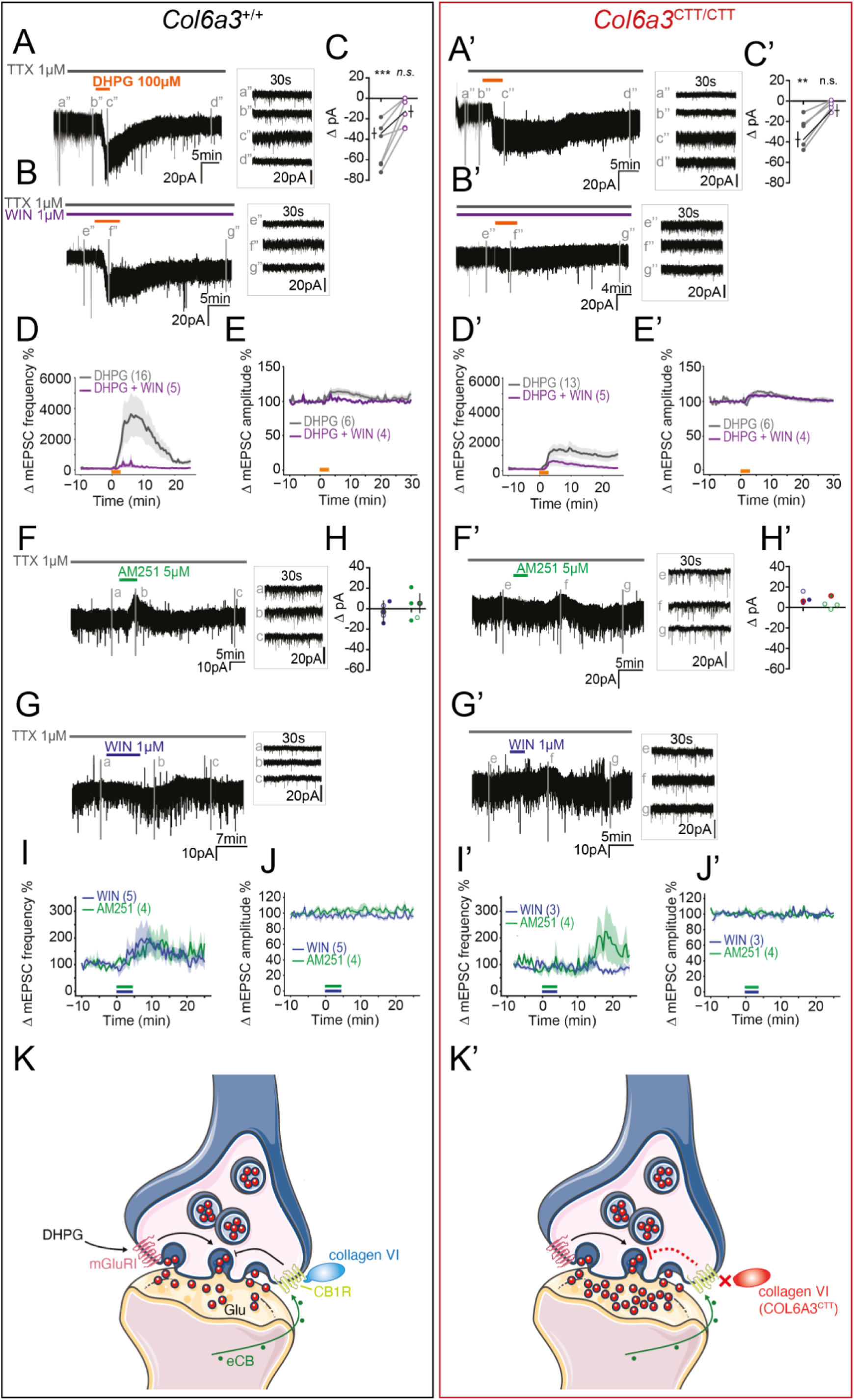
Impaired synaptic homeostasis of *Col6a3*^CTT/CTT^ BPN neurons and improvement with CB1R agonist. (**A,A’**) Representative voltage-clamp response to mGluRI agonist DHPG (orange bar; 100 µM) in wildtype and *Col6a3*^CTT/CTT^ cells respectively in the presence of tetrodotoxin (TTX; 1 µM). (**B,B’**) Representative voltage-clamp response to DHPG in the presence of CB1R agonist WIN 55,212-2 (WIN; purple bar; 1 µM) in wildtype and *Col6a3*^CTT/CTT^ cells respectively in the presence of TTX. (**C,C’**) Quantitative summary of DHPG-evoked current in TTX alone (right) and with WIN (right) in wildtype and *Col6a3*^CTT/CTT^ cells respectively. (**D,D’**) Effect of DHPG on mEPSC frequency in wildtype and *Col6a3*^CTT/CTT^ cells respectively. DHPG causes a large increase in mEPSC frequency which then returns to baseline. This decay effect is impaired in *Col6a3*^CTT/CTT^ mice. The effect of DHPG is blocked by WIN, albeit more effectively in wildtypes (purple traces). Mean and 95% confidence intervals are shown. (**E,E’**) Effect of DHPG on mEPSC amplitude in wildtype and *Col6a3*^CTT/CTT^ cells respectively. The effect of DHPG is blocked by WIN in wildtypes but not in *Col6a3*^CTT/CTT^ mice. Mean and 95% confidence intervals are shown. (**F,F’**) Representative voltage-clamp trace to assess CB1R tone. The effect of CB1R antagonist AM251 (green bar; 5 µM) in the presence of TTX on wildtype and *Col6a3*^CTT/CTT^ cells respectively. (**G,G’**) Representative voltage-clamp traces of CB1R agonist WIN (navy bar; 1 µM) in TTX on wildtype and *Col6a3*^CTT/CTT^ cells respectively. (**H,H’**) Quantitative summary of AM251 and WIN currents (left, WIN, right, AM251). Circles with bold grey (wildtype)/red (CTT/CTT) border are mean ± SEM. Filled circles are statistically significant effects, empty circles were non-significant currents. (**I,I’**) Effect of WIN and AM251 on mEPSC frequency in wildtype and *Col6a3*^CTT/CTT^ cells respectively. Mean and 95% confidence intervals are shown. (**J,J’**) Effect of WIN and AM251 on mEPSC amplitude in wildtype and *Col6a3*^CTT/CTT^ cells respectively. Mean and 95% confidence intervals are shown. (**K,K’**) Schematic of afferent BPN synapses in wildtype and *Col6a3*^CTT/CTT^ mice respectively. DHPG activates mGluRI, stimulating synaptic glutamate release. Postsynaptic neurons produce eCBs which signal retrogradely at presynaptic CB1R, suppressing glutamate release. Collagen VI facilitates CB1R function via the COL6A3 CTD. In *Col6a3*^CTT/CTT^ mice, collagen VI modulation of CB1R is perturbed, disrupting inhibition of glutamate release and resulting in excessive glutamate release. Graphics adapted from Servier Medical Art under a Creative Commons licence (CC BY 3.0).

To assess whether the presumed action of WIN was reflected by changes in glutamate input, we measured mEPSC activity. Unexpectedly, DHPG application potentiated glutamatergic input to BPN cells (WT: baseline 0.83 ± 0.21 Hz versus +DHPG 11.12 ± 3.04 Hz, n=16, RM-ANOVA p = 0.003; *Col6a3*^CTT/CTT^: baseline 2.55 ± 1.22 Hz versus +DHPG 17.81 ± 3.56 Hz, n=13, RM-ANOVA p = 0.0002; Figure 6D, 6D’, 6E, and 6E’). This is in contrast to the previously reported rapid depression of glutamatergic input DHPG causes in accumbal and striatal cells (Kreitzer and Malenka, 2007; Robbe et al., 2002). The change in mEPSC frequency was depressed in *Col6a3*^CTT/CTT^ cells yet the response was prolonged (WT_wash_: 3.03 ± 0.95 Hz; *Col6a3*^CTT/CTT^_wash_: 10.72 ± 2.50 Hz;% decay from t=7 min to 25 min, WT: 68 ± 7%; *Col6a3*^CTT/CTT^: 33 ± 13%, p < 0.03; Figure 6D and 6D’), indicating an overall exaggerated potentiation and thus an impaired adaptation to excitation.

WIN blocked this DHPG-mediated excitation in both wildtype (p=0.12, n=5) and *Col6a3*^CTT/CTT^ cells (p=0.15, n=5), though the effect of WIN on DHPG-induced mEPSC increase was attenuated in *Col6a3*^CTT/CTT^ cells (Figure 6D and 6D’). DHPG also caused an increase in the amplitude of excitatory inputs in a proportion of cells within both genotypes (WT: baseline 10.89 ± 1.19 pA versus +DHPG 12.27 ± 1.14 pA, n= 6 of 15, RM-ANOVA p = 0.02; and *Col6a3*^CTT/CTT^: baseline 10.20 ± 1.27 pA versus +DHPG 11.80 ± 1.30 pA, n= 6 of 14, RM-ANOVA p = 0.02; Figure 6E and 6E’). The remaining cells did not show any statistically significant deviation of amplitude with pharmacological treatment (p = 0.11 and p = 0.09, for WT and *Col6a3*^CTT/CTT^ respectively). When repeated in the presence of WIN, this increase in amplitude in some cells persisted (WT: WIN 11.99 ± 0.76 pA versus +WIN/DHPG 12.77 ± 0.70 pA, n= 4 of 6, RM-ANOVA p = 0.046; and *Col6a3*^CTT/CTT^:WIN 10.55 ± 1.40 pA versus +DHPG 11.43 ± 1.52 pA, n= 4 of 5, RM-ANOVA p = 0.02; Figure 6E and 6E’). WIN did block a DHPG-induced amplitude increase in 2 of the 4 WT cells but none from the *Col6a3*^CTT/CTT^ population. These data suggest a DHPG postsynaptic effect which may be independent of mGluRI, and which is more prevalent in the *Col6a3*^CTT/CTT^ cells. Taken together, these results demonstrate an impaired, though not completely abrogated, effect of CB1R agonism to counteract glutamatergic excitation in *Col6a3*^CTT/CTT^ cells.

To probe the endogenous eCB tone, we treated cells with either the CB1R inverse agonist AM251 (5µM; green bar, Figure 6F and 6F’) or with the CB1R agonist WIN (1µM; navy bar, Figure 6G and 6G’) in TTX. AM251 preferentially evoked an outward current in WT cells (n= 2 of 3 cells with significant current; overall average ΔI of +4.95 ± 9.37 pA; Figure 6H) while WIN caused mixed currents in only half the cells tested (n= 2 of 4 cells with significant current; overall average ΔI of −3.40 ± 10.73 pA). In contrast, AM251 evoked an outward current in only 1 cell from *Col6a3*^CTT/CTT^ BPN slices (I of 11.39 pA, n= 1 of 4; Figure 6H’) while WIN evoked a significant outward current in 2 of 3 cells examined (overall average ΔI of +6.09 ± 1.56 pA). However, compared to the group baseline values in either genotype, neither compound reached statistical significance, suggesting little endogenous eCB tone on the postsynaptic membrane.

When we assessed presynaptic eCB tone, AM251 and WIN both tended to increase mEPSC activity in WT cells, yet the effect did not reach significance (WT: baseline 3.07 ± 1.31 Hz versus +AM251 6.53 ± 2.6 Hz, n= 4, RM-ANOVA p = 0.15; and WT: baseline 1.62 ± 0.68 Hz versus +WIN 3.04 ± 1.34 Hz, n= 5, RM-ANOVA p = 0.17; Figure 6I). There was no effect on mEPSC frequency in the *Col6a3*^CTT/CTT^ cells (baseline 3.56 ± 1.49 Hz versus +AM251 4.51 ± 1.53 Hz, n= 4, RM-ANOVA p = 0.35; and *Col6a3*^CTT/CTT^: baseline 2.30 ± 1.32 Hz versus +WIN 1.63 ± 0.36 Hz, n= 3, RM-ANOVA p = 0.38; Figure 6I’) nor on mEPSC amplitude in either genotype (Figure 6J and 6J’). These data suggest there is little to no endogenous eCB tone in the BPN prior to DHPG-stimulation. Collectively, the data demonstrate impaired synaptic homeostasis in *Col6a3*^CTT/CTT^ neurons, apparently due to impaired CB1R function (Figure 6K and 6K’).

### CB1R agonist rapidly improves motor performance in *Col6a3*^CTT/CTT^ mice

We showed that BPN neurons of *Col6a3*^CTT/CTT^ mice were impaired in their ability to restore homeostasis to excitatory input signaling, and hypothesized that this mediates at least part of their dyskinetic phenotype. We also showed that WIN could still restrain excitatory input potentiation in the BPN in *Col6a3*^CTT/CTT^ mice, albeit with reduced efficacy. This led us to hypothesize that CB1R agonism may be an effective treatment for the dyskinesia seen in *Col6a3*^CTT/CTT^ mice. We subjected mice to a randomized crossover study testing the effect of acute CB1R agonism on motor performance. We found that WIN treatment 2 hours before the beam walk test (Morgese et al., 2007) significantly improved hindlimb coordination (Figure 7 and S7). Applying stratified analysis to evaluate the effect of WIN on values above the population median, we observed significant improvements in *Col6a3*^CTT/CTT^ mice in hindlimb coordination and traversal time on the 12 and 15 mm beams (paired *t*-test: hindlimb coordination, 12 mm beam: p = 0.03, t = 2.3, df = 16; 15 mm beam: p = 0.01, t = 2.7, df = 16; traversal time, 12 mm beam: p = 0.01, t = 1.5, df = 17; 15 mm beam: p = 0.03, t = 2.4, df = 17), as well as number of slips on the 20 mm beam (paired *t*-test, p = 0.02, t = 2.9, df = 9).

**Figure 7.**
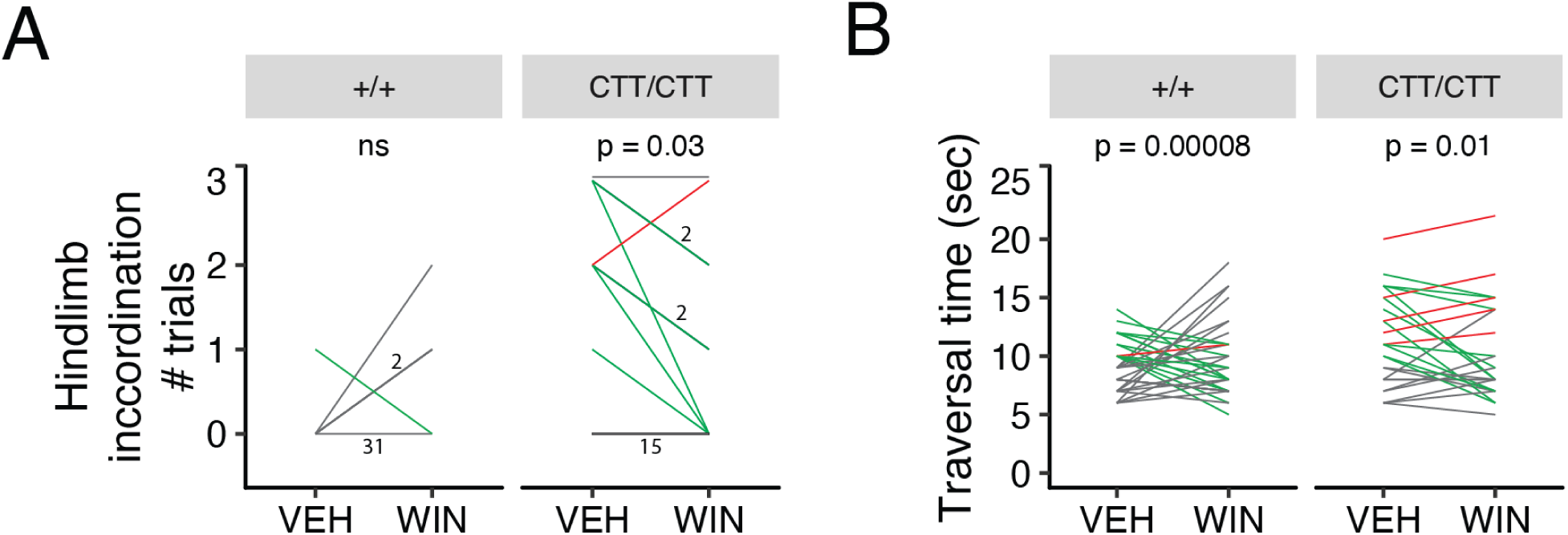
Cannabinoid-mediated improvement of motor control in *Col6a3*^CTT/CTT^ mice. (**A**) Effect of WIN treatment in randomized crossover design study on hindlimb incoordination on 12 mm rectangular beam. Green, performance improved by WIN; red, performance worsened with WIN; grey, baseline value of 0 (unimpaired) or performance unchanged with WIN. Number indicates number of mice with indicated trajectory. (**B**) Effect of WIN treatment on traversal time of 12 mm rectangular beam. Green, performance improved by WIN; red, performance worsened with WIN; grey, baseline value below median (9.5 sec) or performance unchanged with WIN. WIN was particularly beneficial for mice with poor baseline performance. Male and female mice aged 12-28 wk, n=26-34 per genotype; same mice as shown in Figure 2C and 2D. See also Figure 7-1.

## DISCUSSION

We establish a molecular and synaptic mechanism through which the unique C-terminal domain of collagen α3 (VI) regulates motor function, and through which variants affecting this domain engender the movement disorder dystonia. The mechanism implicates the cannabinoid receptor CB1R, which can be targeted noninvasively to improve motor performance.

### *Col6a3*^CTT^ mouse model

The *Col6a3*^CTT^ mutation caused substantial NMD of the transcript, meaning that *Col6a3*^CTT/CTT^ expressed a truncated COL6A3 protein at significantly lower levels than wildtypes. Nevertheless, the mice had a dyskinetic phenotype which was not occluded by myopathy, unlike mice with complete loss of COL6A3 function which are overtly myopathic (Pan et al., 2013, 2014). Like patients with variants affecting the COL6A3 C-terminal domain, the dyskinetic phenotype of *Col6a3*^CTT^ mice was recessive, with heterozygous mice showing normal motor performance (Figure 2). *Col6a3*^CTT/CTT^ mice also show modestly reduced muscle strength in older mice. As reduced muscle strength is seen in unrelated mouse models of dystonia such as a cholinergic system torsinA-deficient mouse (Pappas et al., 2018), it is unclear whether it is in this case related to an effect of the CTT mutation on collagen VI function in muscle. Mitigating against this possibility is the observation that muscle histology is normal in young *Col6a3*^CTT/CTT^ mice, and even aged *Col6a3*^CTT/CTT^ have only a moderate myopathy (Pan et al., 2013, 2014).

### The collagen-cannabinoid connection

Collagen VI is deposited in a wide range of tissues, including skeletal muscle, adipose tissue, cartilage, skin, and nervous system (Cescon et al., 2015). Interactions with a variety of extracellular matrix components and cell surface receptors have been shown, including other collagens, glycoproteins, and integrins (Cescon et al., 2015). The functional repertoire extends well beyond structural functions to include cytoprotection, regulation of myelination, macrophage recruitment, and hair growth, yet in each of these cases the mechanism, and particularly the interaction partner, remains largely unknown (Cescon et al., 2015). Here, we identify a specific receptor through which collagen VI can modulate cellular and circuit functions. CB1R has a highly plastic orthosteric binding pocket, enabling it to respond to a variety of ligands of considerably different shapes and sizes (Hua et al., 2017). Thus, collagen VI may act as a positive allosteric modulator of CB1R.

### Basal pontine plasticity

The BPN links the cerebral cortex and cerebellum in an integral motor control circuit, which mediates the selection and denoising of cortical representations during motor learning (Guo et al., 2019; Wagner et al., 2019). We now show that BPN neurons are indeed capable of input potentiation, which is homeostatically counteracted by eCBs. As eCB retrograde inhibition is operative in the BPN, it may contribute to some of the key features of adaptive motor control, such as surround inhibition, which appear disrupted in dystonia. We did not find evidence for eCB-mediated synaptic depression, as is observed in the striatum and accumbens (Kreitzer and Malenka, 2007; Robbe et al., 2002). However, other paradigms for evoking synaptic depression, such as high frequency afferent stimulation, would be an interesting avenue of future investigation.

Other studies have associated collagen VI with neuroprotection. Specifically, disruption of collagen VI increased apoptosis, while exogenous collagen VI reduced stress-induced apoptosis in primary cortical/hippocampal neuronal cultures (Cescon et al., 2016; Cheng et al., 2009). We did not detect robust evidence of apoptosis in *Col6a3*^CTT/CTT^ mice under basal conditions (data not shown). Further studies are required to determine whether challenging the circuit, e.g. with physiological stress, may induce apoptosis. Intriguingly, cannabinoid signaling is protective against excitotoxicity (Marsicano et al., 2003). Further studies are required to clarify whether this mechanism is at least partially responsible for the neuroprotective effect of collagen VI.

### Cannabinoid modulation of motor performance

We demonstrate that cannabinoid augmentation with a CB1R agonist can improve motor outcomes in impaired *Col6a3*^CTT/CTT^ mice. Though COL6A3-related dystonia is rare, our results suggest that cannabinoid augmentation may have therapeutic potential in these patients. Furthermore, this potential may generalize to other forms of dyskinesia. Indeed, several case studies and small open-label studies report a beneficial effect of cannabinoid treatment in dyskinesia (Koppel, 2015). Our mechanistic findings reported here provide a firm evidentiary basis and give fresh impetus to investigating cannabinoid-related treatment of dyskinesias.

## ACKNOWLEDGEMENTS

We thank Irmgard Zaus, Julia Vandrey, Monika Taschner, and Celestine Dutta for technical assistance, Raimund Wagener for antibodies, and Feng Zhang for plasmids. This work was supported by the following grants: Deutsche Forschungsgemeinschaft LA 3830/1-1 to D.D.L and WI 1820/8-1 to J.W.; European Research Council Starting Grant iNAPS to R.H.W.; Deutscher Akademischer Austauschdienst Rückkehrstipendium to D.D.L.; Canadian Institutes of Health Research (Foundation Grant FDN-167281), the Canadian Institutes of Health Research and Muscular Dystrophy Canada (Network Catalyst Grant for NMD4C), the Canada Foundation for Innovation (CFI-JELF 38412), and the Canada Research Chairs program (Canada Research Chair in Neuromuscular Genomics and Health, 950-232279) to H.L.

## AUTHOR CONTRIBUTIONS

Conceptualization, D.D.L, M.Z., and J.W; Funding acquisition, D.D.L, R.H.W., and J.W.; Investigation, D.D.L, R.H.W., E.L., K.T., G.H., N.L.S., J.M-P., A.V.S., D.L., S.S., M.J.P., and H.L.; Project administration, D.D.L. and J.W.; Resources, R.H.W., G.H., M.W., and J.W.; Supervision, D.D.L, A.G., S.M.H., M.S., M.W., and J.W.; Visualization, D.D.L. and R.H.W.; Writing – original draft, D.D.L; Writing - review & editing, D.D.L, R.H.W., M.Z., S.M.H., M.W., and J.W.

## MATERIALS AND METHODS

### Experimental Model and Subject Details

159 mutant or wild-type mice on a C57BL/6 background aged 0 to 402 days were used in the experiments. 6 male wildtype and 3 male Col6a3^CTT/CTT^ mice were used for *in situ* hybridization studies. 51 wildtype, 16 Col6a3^+/CTT,^ and 36 Col6a3^CTT/CTT^ mice were used for behavioural studies. 6 male wildtype and 6 male Col6a3^CTT/CTT^ mice were used for muscle histology studies. 2 male and 2 female wildtype mice were used to prepare brain lysates for binding partner studies. 15 wildtype and 11 Col6a3^CTT/CTT^ mice were used for electrophysiology studies. 5 wildtype mice were used to prepare primary cortical neurons. Sample size for behavioural experiments was in accordance with guidelines set by the Stanford Behavioral and Functional Neuroscience Laboratory, which conducted the initial phenotypic analysis. Sample size for electrophysiology was determined based on previously published studies, and statistical significance was calculated using post hoc tests. For the crossover study, treatment order was randomized with the ‘randomizr’ R package, complete_ra function. Experimenters were blinded to animal genotype in all behavioural studies. All animal work was performed in accordance with the German Animal Welfare Act, European Union Directive 2010/63, and the National Institutes of Health Guide for the Care and Use of Laboratory Animals, and approved by the Government of Upper Bavaria and the Stanford Administrative Panel on Laboratory Animal Care.

Animals were kept in individually ventilated Tecniplast Greenline GM 500 cages (501 cm² floor area) with laboratory animal bedding (wood fibre/wood chips) and autoclaved nesting material (nestlets, cardboard houses, plastic igloos, or cellulose). Animals were maintained under specified pathogen-free (SPF) conditions in fully air-conditioned rooms with a 12/12 hr light/dark cycle, a temperature of 20-24°C and humidity of 45-65%. The cages were changed on average once a week, and animals received sterile filtered water and standard rodent diet *ad libitum*.

### Histology

For *in situ* hybridization, animals were euthanized by cervical dislocation. Fresh brains were extracted, snap frozen in −60°C isopentane (2-methylbutane), sectioned at 30 µm thickness on a cryostat, and collected on Superfrost slides. For chromogenic *in situ* hybridization, sections were fixed in 4% paraformaldehyde in phosphate-buffered saline (PBS) for 10 min, then rinsed in PBS. After three 10 min PBS washes, sections were acetylated in 1.35% triethanolamine, 0.02 N hydrochloric acid, and 0.25% (v/v) acetic anhydride for 10 min. Next, sections were permeabilized with 1% (v/v) Triton X-100 in PBS, followed by three 10 minute PBS washes. Sections were then prehybridized with hybridization solution (50% formamide, 10% dextran sulfate, 2% Denhardt’s solution, 1 mg/ml yeast transfer RNA, 300 mM NaCl, 10 mM Tris-HCl pH 6.5, 5 mM EDTA pH 8, 50 mM NaH_2_PO_4_, 50 mM Na_2_HPO_4_) for 2-4 hrs.

To synthesize the riboprobe, a 509 bp cDNA fragment spanning the C-terminal encoding portion of the transcript (base 9,548-10,056 of Ensembl transcript ENSMUST00000056925.15, spanning exons 40-43) was cloned from mouse brain cDNA into pCRII-TOPO (Thermo Fisher). The plasmid was linearized and subjected to *in vitro* transcription with digoxigenin labeling (DIG RNA labeling kit, Sigma). Sections were hybridised with 500 ng/ml riboprobe in hybridization solution at 72°C overnight. Next, sections were incubated with 0.2X saline sodium citrate (SSC; 30 mM NaCl, 3 mM trisodium citrate, pH 7) at 72°C for 45 min, then washed in 0.2X SSC for 5 min. To detect digoxigenin, sections were blocked in B1 (100 mM Tris-HCl pH 7.5, 150 mM NaCl, 0.1% Triton X-100) with 1% heat-inactivated goat serum for 1 hr, then incubated with anti-digoxigenin antibody coupled to alkaline phosphatase (Sigma 11093274910, 1:3,500) in the same solution overnight. Sections were then washed (10 min/wash) three times in B1 and twice in B2 (100 mM Tris-HCl pH 9, 100 mM NaCl, 50 mM MgCl_2_). Sections were then incubated in B2 with 2% nitro blue tetrazolium/5-bromo-4-chloro-3-indolyl-phosphate (NBT/BCIP, Sigma) until adequate colour development. The reaction was stopped with B1, and slides were coverslipped with polyvinyl alcohol.

For fluorescent *in situ* hybridization (FISH), sections were fixed in 4% paraformaldehyde in PBS at 4°C for 1 hr, then rinsed twice with PBS. Sections were then dehydrated in ascending concentrations of ethanol (50%, 70%, 100%, 100%; 5 min each). Slides were baked at 37°C for 30 min, then sections were demarcated with a hydrophobic barrier pen. Sections were pretreated with RNAscope Protease IV Reagent (ACD) for 30 min and rinsed twice in PBS. Probes (ACD) were added to sections (C1: *Slc17a7*, C2: *Col6a3*, C3: *Dagla*) and hybridized at 40°C for 2 hr. Sections were then exposed to amplification reagents (Amp1-FL, 30 min; Amp2-FL, 15 min; Amp3-FL, 30 min; Alt4-FL Alt B, 15 min) at 40°C with two washes of two minutes with RNAscope wash buffer between each step. After 30 sec DAPI exposure, sections were coverslipped with Aqua-Poly/Mount (Polysciences, Inc). Images were obtained with an Axio Scan.Z1 digital slide scanner (Zeiss, Jena, Germany) equipped with a 20x magnification objective. Images were evaluated using the commercially available image analysis software Developer XD 2 (Definiens AG, Munich, Germany). Thresholds were set empirically for each channel and the same thresholds were applied to all images, allowing nuclei to be classified as positive or negative for each transcript.

For hematoxylin and eosin staining, tibialis anterior muscle was dissected, embedded in OCT (Tissue-Tek), snap frozen in isopentane, and sectioned on a cryostat at 12 µm thickness. Sections were fixed with 4% paraformaldehyde in PBS for 5 min, then washed in tap water. Next, sections were incubated in Mayer’s hematoxylin solution for 5 min, washed with tap water, incubated with 0.5% eosin and 1% acetic acid for 10 min, and washed three times with distilled water. Sections were then dehydrated (70% ethanol for 1 min, 90% ethanol for 30 sec, 100% ethanol for 30 sec, xylene for 30 sec) and coverslipped with Depex. Images of stained muscle sections were captured on a Axio Scan.Z1 digital slide scanner equipped with a 20x magnification objective, and analysed with FIJI(Schindelin et al., 2012). Transverse fibres were traced using a freehand selection tool by an investigator blinded to the genotype, and measurements of the fibre size (μm^2^) and circularity (0-1, with 1=perfect circle and 0.2=very angular) were recorded.

### Genetic engineering

To generate *Col6a3*^CTT^ mice, we linearized the pSpCas9(BB)-2A-Puro (PX459) plasmid (a gift from Feng Zhang; Addgene plasmid # 48139) with BbsI and dephosphorylated it with calf intestinal alkaline phosphatase. Guide RNA (gRNA) oligonucleotides (5’-CACCGCACCTCTCGCATCCGGCTGA-3’ and 5’-AAACTCAGCCGGATGCGAGAGGTG C-3’) were simultaneously phosphorylated and annealed as follows: 100 pmol of each oligonucleotide were mixed with T4 ligase buffer and T4 polynucleotide kinase (5 U), and incubated at 37°C for 30 min, 95°C for 5 min, and then cooled to 25°C at a rate of 0.1°C/s. Annealed and phosphorylated oligonucleotides were then ligated into linearized and dephosphorylated PX459 with T4 ligase. Ligation product was transformed into competent bacterial cells, and bacterial clones with successful integration of the gRNA were used for plasmid cloning.

Purified plasmid was transfected into V6.5 mouse embryonic stem (ES) cells with Fugene HD (370 ng plasmid, 100,000 cells). One day after transfection, successfully transfected cells were selected with puromycin (2 µg/ml) for 2 days. Surviving cells were clonally diluted, expanded, and picked for genotyping with the screening primers 5’-AGTGCCCTGTATTCCCAACA-3’ and 5’-AAAGCGTTGATGAGCTGTCG-3’.

In one clone, 5’-CAACTGTCCGCGG-3’ was deleted and replaced by 5’-CCCACAAAGAGAGAGAGTCA-3’, yielding a frameshift and premature stop codon in exon 36. In the predicted protein product, the 15 amino acid sequence NCPRGARVAVVTYNN is replaced by PQRERVRCPRGCGHL followed by the stop codon, resulting in severe disruption of the C1 domain (out of 180 amino acids, 36 intact, 15 altered, and 129 missing) and complete absence of the C2-C5 domains.

Altered ES cells were injected into blastocysts, which were transferred to the uterus of pseudopregnant female mice. Chimeric offspring were backcrossed to C57/BL6J mice to obtain germline transmission. Mice were genotyped with the primers 5’-AGAGAGCAACTGTCCGCG-3’ (wildtype-specific sense primer), 5’-AGCCCACAAAGAGAGAGAGT-3’ (CTT-specific sense primer), and 5’-AAAGCGTTGATGAGCTGTC-3’ (common antisense primer) in two separate PCRs (wildtype sense and common antisense primers, yielding a 360 bp product, and CTT-specific sense and common antisense primers, yielding a 367 bp product). The sequence of the CTT allele was verified by RNA-seq (Figure S1A).

We used CRISPOR (Concordet and Haeussler, 2018) to identify potential off-target sites for the guide RNA used. No sites had below 4 mismatches with the guide sequence. 11 exonic sites (out of 44 total) had 4 mismatches. None of the 44 were PAM-adjacent. Of the 11 exonic sites, 9 were covered by at least 1 BPN RNA-seq read extending at least 15 bp in either direction from the PAM. Within these reads, there were no sequence variants.

### Quantitative PCR and RNA-seq

Muscle and brain regions were dissected and homogenized (muscle was frozen on dry ice and crushed with a pestle and mortar; brain tissue was homogenized with a syringe and 20 gauge needle in RLT buffer). RNA was extracted with the RNeasy kit (Qiagen) and reverse transcribed with GoScript Reverse Transcriptase with random primers (Promega). Real-time quantitative PCR was done using FAM-MGB-labeled *Col6a3* (Mm00711678_m1) and *Polr2a* (Mm00839502_m1) TaqMan assays.

For RNA-seq, RNA was prepared as for qPCR. RNA integrity number (RIN) was determined with the Agilent 2100 BioAnalyzer (RNA 6000 Nano Kit, Agilent). For library preparation, 1 μg of RNA was poly(A) selected, fragmented and reverse transcribed with the Elute, Prime and Fragment Mix (Illumina). End repair, A-tailing, adaptor ligation and library enrichment were performed as described in the Low Throughput protocol of the TruSeq RNA Sample Prep Guide (Illumina). RNA libraries were assessed for quality and quantity with the Agilent 2100 BioAnalyzer and the Quant-iT PicoGreen dsDNA Assay Kit (Life Technologies). RNA libraries were sequenced as 100 bp paired-end runs on an Illumina HiSeq2500 platform. RNA-seq reads were demultiplexed and mapped with STAR(Dobin et al., 2013) (version 2.4.2a) to the mm9 genome assembly (UCSC Genome Browser build) using default parameters, except: min length of chimeric segment: 20; quantMode GeneCounts (to quantify reads per gene); 2 pass mapping; unmapped reads kept; all sam attributes output.

### Preparation of protein lysates

Tibialis anterior muscles were dissected, frozen on dry ice, and pulverised with a pestle and mortar. Mouse brain lysates were prepared from fresh microdissected wildtype brain tissue. “LSN+” (Figure 3B and Figure 3D) included also medial septal nucleus and striatum. Dissected tissue was placed in ice cold lysis buffer (150 mM NaCl, 1% NP-40, 50 mM Tris-HCl pH 8, and cOmplete Mini Protease Inhibitor, Merck) and homogenized with a syringe and 20G needle. The sample was agitated at 4°C for 30 min, then centrifuged at 10,000 rpm for 10 min at 4°C. The supernatant was used for subsequent experiments, with quantification by the DC Protein Assay (Bio-Rad).

### Western blots

10 µg of tissue lysate was mixed with Lämmli buffer, incubated at 95°C for 5 min, and electrophoresed (SDS-PAGE, 125V) on 4-20% polyacrylamide gels. Proteins were transferred to nitrocellulose membranes by wet blot, blocked in 5% milk in Tris-buffered saline with 1% Tween-20 (TBST) for 1 hr, and incubated with primary antibodies (mouse anti-vinculin V9131, Sigma; and from Heumüller et al., 2019, rabbit anti-COL6A3-N 1:500) in TBST with 5% milk at 4°C overnight. Membranes were washed 3 times in TBST, then incubated with secondary antibody (HRP-conjugated donkey anti-rabbit 406401, BioLegend, 1:1000; HRP-conjugated goat anti-mouse Cay10004302-1, Cayman, 1:1000). Membranes were again washed 3 times in TBST, and HRP was chemiluminescently detected with ECL Prime Detection Reagent (Amersham) on a Fusion Pulse imager (Vilber).

### Mouse behavioural tests

For the beam walk task, mice were trained to traverse a beam (square profile, 20 mm diameter, length 90 cm, 15 cm above the surface) with the home cage on one end under normal light. Mice were guided to the home cage if necessary. Mice completed 3 successive traversals in training. Experiments commenced approximately 1 week after training. Some mice received injections prior to testing. The injections were administered in the testing room between 08:30 and 10:30. Vehicle (0.33% DMSO) or the CB1R agonist WIN 55,212-2 (WIN; 1 mg/kg; Cayman Chemical) were injected intraperitoneally (10 µl/g body weight). Mice in the treatment study received both WIN and vehicle in a randomized crossover design, with the two conditions separated by 1-8 weeks; see previous section). Testing commenced 2 hours after injection.

In testing, performed under normal light between 10:30 and 12:30, mice traversed each of 4 beams 3 times each consecutively: 20 mm square profile, 22 mm circular profile, 12 mm square profile, and 15 mm circular profile). The traversal time, number of slips and falls, and the presence of hindlimb incoordination were recorded. Hindlimb coordination was characterized by an apparent inability to coordinate the hindlimbs towards forward locomotion (see Movie S1). The effects of WIN treatment were evaluated separately in mice with baseline values above the overall median for each parameter (impaired mice).

For the grip strength test, maximum grip strength of forelimbs and combined forelimbs and hindlimbs was measured. The mice were allowed to grasp a grid connected to a force sensor (Bioseb, Chaville, France) while being pulled away. Three consecutive trials were carried out for forelimbs and all limbs, and the means of the respective trials were used for analysis.

The rotarod test is designed to evaluate the motor coordination and balance of rodents by forcing the subjects to run. The subjects are placed on a rotating rod (Med Associate Inc, Model ENV-575M) with a steady acceleration and the latency to fall is recorded. The subjects fall safely 16.5cm below the rotating rod. The experiment consists of one day of training and one day of testing. On the training day, subjects are placed on the rod accelerating from 2-20rpm for 5 minutes until they fall or the 5 minutes trial is complete. This process is repeated for a total of 3 trials with 15 minutes Inter-trial-interval (ITI). The subjects receive a resting day after the training day. On the testing day, the mice are tested with constant speed of 16rpm for a total of 5 trials with 15 minutes ITI. The rod is cleaned with 1% Virkon between trials.

The Activity Chamber is a simple assessment test used to determine general activity levels, gross locomotor activity, and exploration habits in rodents. Assessment takes place in a square arena, 43.2×43.2 cm, mounted with three planes of infrared detectors, within a specially designed sound attenuating chamber, 66×55.9×55.9 cm. The animal is placed in the center of the testing arena and allowed to freely move for 10 minutes in the dark (with red lighting) while being tracked by an automated tracking system. Distance moved, velocity, jump counts, vertical counts and resting time in the arena are recorded. At the conclusion of each trial the surface of the arena is cleaned with 1% Vikron.

### Synthesis of tagged proteins

Mouse COL6A3 sequence without 48 amino acid cleaved leader sequence was amplified by PCR from mouse brain cDNA and cloned into the transfer vector pFastBac-EGT-N (Helmholtz Center Munich) in frame with the N-terminal secretion signal ecdysteroid UDP-glucosyltransferase (EGT), 6xHis tag, and Flag tag. C4 domain sequence was removed with the Q5 site-directed mutagenesis kit (NEB). Mouse prolyl 4-hydroxylase alpha- and beta-chain (P4HA1 and P4HB) sequences were amplified by PCR from mouse brain cDNA and cloned into the bicistronic transfer vector pFastBac-Dual (ThermoFisher). Constructs were validated by sequencing prior to site-specific recombination with the baculovirus genome provided by its host E.coli strain DH10Bac (Bac-to-Bac-System, Invitrogen). Positive recombinant clones were identified by blue-white screening and further confirmed by PCR. Recombinant viral DNA was extracted under alkaline conditions, followed by ethanol precipitation (Birnboim and Doly, 1979) and used for transfection into insect cells (Fugene HD, Promega). Baculovirus recombinant for COL6A3^WT^ or COL6A3^ΔC4^ were propagated in Sf21 cells and supernatant (SN) was kept at 4°C for further analysis and protein production. Low passage viral titer was determined by quantitative PCR (Lo and Chao, 2004). Briefly, viral DNA was purified from SN (High Pure Viral Nucleic Acid Kit, Roche) and PCR was set up with Brilliant II qPCR Low ROX Master Mix (Agilent Technologies). qPCR reaction and data were acquired on an Agilent Mx3005P qPCR System.

COL6A3^WT^ and COL6A3^ΔC4^ were produced in High Five insect cells. For protein production, High Five cells were seeded 1 day prior to infection at 1×10^6^ cells/mL in InsectXpress medium (Lonza). Cells were co-infected with low-passage recombinant baculovirus for COL6A3^WT^ or COL6A3^ΔC4^ along with P4HA1/P4HB at a MOI of 2 and 0.2 respectively, in the presence of 80 µg/mL ascorbic acid (AppliChem). Cells were incubated at 21°C on an orbital shaker. SN was harvested 3 days post infection by centrifugation, adjusted to final concentrations of 20 mM Tris pH 7.1 and 0.02% thioglycerol (TG), and filtered (0.45µm Stericup, Merck Millipore). Collagens were captured by ion exchange chromatography using a CaptoS-5mL prepackaged column (GE Healthcare) and eluted with 20 mM Tris pH 7.1, 0.02% TG, and 250 mM NaCl. Positive fractions were pooled, concentrated (Amicon, 30kD MWCO, Merck Millipore) and polished by gel filtration using an S200pg (GE Healthcare) buffered in 20 mM Tris pH 7.1, 0.02% TG, and 300 mM NaCl. Positive fraction were pooled, concentrated and lyophilized in aliquots. Purity was determined by SDS-PAGE and Coomassie staining (Page Blue, Fermentas), and protein concentration was determined photometrically. Purified tagged COL6A3 proteins are depleted, but plasmids to produce them are freely available from the authors.

### Immunoprecipitation

For identification of COL6A3 binding partners, anti-FLAG M2 magnetic beads (Merck) were used; 20 µl beads were diluted in 100 µl Tris-buffered saline (TBS) for each immunoprecipitation reaction. The beads were washed three times with TBS, then incubated with an empirically optimized 4 µg of FLAG-tagged COL6A3 protein in TBS with agitation for 1 hr at room temperature. The beads were again washed three times with TBS. Bait-bound beads were then incubated with an empirically optimized 250 µg of brain lysate in a volume of 1 ml with agitation at room temperature for 2 hr. The beads were again washed three times with TBS. Bait-bound proteins were eluted from the beads with Lämmli buffer (80 mM Tris-HCl pH 6.8, 2% SDS, 10% glycerol, 5% ß-mercaptoethanol, and 0.02% bromophenol blue) at 95°C for 5 min.

### Sample preparation for liquid chromatography-mass spectrometry analysis

For identification of COL6A3 binding partners, Lämmli eluates were subjected to a modified filter-aided sample preparation (FASP) procedure as described (Grosche et al., 2016; Wiśniewski et al., 2009). Briefly, proteins were diluted in ammonium bicarbonate prior to reduction and alkylation using dithiothreitol and iodoacetamide. Urea was added to a final concentration of 4 M, followed by centrifugation of denatured proteins on a filter device (Sartorius, Vivacon 500). After several washing steps with 8 M urea and ammonium bicarbonate, proteins were subjected to on-filter proteolysis with Lys-C (Wako Chemicals) and trypsin (Promega). Tryptic peptides were eluted by centrifugation and acidified with trifluoroacetic acid prior to liquid chromatography-mass spectrometry (LC-MSMS) analysis. For whole proteome analysis of wildtype vs. *Col6a3*^CTT/CTT^ brain tissue, equal amounts of ground brain tissue were lysed in 8 M urea in 0.1 M Tris-HCl pH 8.5 using the Precellys 24 tissue homogenizer (Bertin Instruments). Crude lysates were subjected to FASP digest as described above.

### Quantitative LC-MSMS analysis

Digested IP samples were measured in data-dependent acquisition mode on a Q-Exactive HF mass spectrometer (Thermo Fisher) online coupled to an Ultimate 3000 RSLC (Dionex, part of Thermo Fisher). Peptides were trapped on a C18 pre-column cartridge prior to separation on a nanoEase MZ HSS T3 Column (100Å, 1.8 µm, 75 µm x 250 mm, Waters) in a non-linear acetonitrile gradient over 95 minutes. Both the precursor (scan range 300 – 1500 m/z) and TOP10 MS2 spectra of charges 2-7 were acquired in the orbitrap mass detector of the mass spectrometer, at resolutions of 60,000 and 15,000 respectively. Maximum injection times were set to 50 ms for MS and MS2 scans and dynamic exclusion to 30 s. Acquired raw files were analyzed in the Progenesis QI software for MS1 intensity based label-free quantification (version 3.0, Nonlinear Dynamics, Waters) as described previously (Frik et al., 2018; Hauck et al., 2010). MSMS spectra were searched against the Swissprot mouse database (16,868 sequences, Release 2017_02) using the Mascot search engine (Matrix Science). Search settings were: enzyme trypsin, 10 ppm peptide mass tolerance and 0.02 Da fragment mass tolerance, one missed cleavage allowed, carbamidomethylation was set as fixed modification; methionine oxidation and asparagine or glutamine deamidation were set as variable modifications. Applying the mascot percolator algorithm (Brosch et al., 2009) resulted in a peptide false discovery rate of 0.49%. The abundances of all unique peptides per protein were summed. Resulting normalized abundances of the individual proteins were used for calculation of fold changes and significance values by Student’s *t*-test.

Whole proteome analysis of wild type vs. *Col6a3*^CTT/CTT^ brain tissue was performed in data-independent acquisition mode on the LC-MSMS setup described above, as described previously (Lepper et al., 2018; Mattugini et al., 2018). MS spectra from 300 to 1650 m/z were recorded at 120000 resolution with an AGC (automatic gain control) target of 3 x 10^6^ and a maximum injection time of 120 ms. The mass range was divided into 37 data-independent acquisition windows of unequal size based on previous data-dependent acquisition runs, with each a scan resolution of 30,000 and an AGC target of 3 x 10^6^. Generated raw files were analysed using the Spectronaut X software (Biognosys; Bruderer et al., 2015) employing an in-house mouse spectral meta-library which was generated using Proteome Discoverer 2.1 (Thermo Fisher), Byonic search engine (Protein Metrics) and the Swissprot mouse database (release 2017_02). Quantification was based on the sum of MS2 area levels of all unique peptides per protein fulfilling the *q* value percentile 0.25 setting with a maximum peptide false discovery rate of 1%. Resulting protein abundances were again used for calculation of fold changes and significance values by Student’s *t*-test.

### Primary mouse cortical neurons

Cortices of newborn mice were dissected and placed in ice cold Hank’s balanced salt solution (HBSS) supplemented with 20% fetal bovine serum, 4.2 mM NaHCO_3_, and 1 mM HEPES. Tissue was washed 3 times in HBSS, then incubated with papain (Neuron Isolation Enzyme, Thermo Fisher) at 37°C for 15 min. Digested tissue was then washed 3 times in HBSS and 3 times in plating medium (Neurobasal-A with GlutaMax, B27, and penicillin-streptomycin). DNase I (0.05 U/µl) was added to the final wash. Tissue was then triturated, filtered through a 40 µm strainer, and plated on poly-D-lysine-coated coverslips at 250,000 cells per cm^2^.

### Wash and bind assay

Mouse primary neurons at 2 DIV were used. COL6A3^WT^ or COL6A3^ΔC4^ were added (1 µg per well of 24 well plate, n=3 replicates per condition) and incubated for 2 hr at 37°C. Cells were then washed 5 times with PBS, fixed with 4% paraformaldehyde for 15 min, and labeled with 6x-His Tag Antibody coupled to Alexa 555 (Thermo Fisher, MA1-21315-A555). Coverslips were mounted on slides (Aqua-Poly/Mount) and images obtained while blinded to condition with a LSM880 confocal microscope (Zeiss, Jena, Germany) equipped with a 20x magnification objective.

### cAMP assay

Mouse primary neurons at 2 DIV were used, and the cAMP-Glo Max Assay (Promega) was used for cAMP detection and quantification. The induction medium consisted of PBS with 500 µM 3-isobutyl-1-methylxanthine (IBMX), 100 µM Ro-20-1724, 10 µM forskolin, and 40 mM MgCl_2_. Cells were incubated with induction medium and test compounds for at 37°C for 15 min. Samples were then analyzed using the manufacturer’s luminescence protocol.

### Single cell RNA-seq analysis

Cluster trinarization scores are as provided by the authors (Zeisel et al, 2018). We used the full set of hindbrain neurons from Zeisel et al, 2018 (“L6_Hindbrain_neurons.loom”). This dataset contained 1,144 cells, of which 97 expressed Col6a3. We performed gene-wise Spearman correlation in R (cor.test) using all 1,144 hindbrain neurons and all 27,998 genes.

### Electrophysiology

We prepared coronal brain slices (200 μm) in ice-cold, oxygenated (95% O_2_, 5% CO_2_) sucrose-based artificial cerebrospinal fluid (aCSF) containing the following (in mM): 250 sucrose, 2.5 KCl, 1.24 NaH_2_PO_4_, 10 MgCl_2_, 10 glucose, 26 NaHCO_3_, 0.5 CaCl_2_ (305 mOsm/L). We then incubated slices in ACSF containing the following (in mM): 124 NaCl, 2.5 KCl, 1.24 NaH_2_PO_4_, 1.3 MgCl_2_, 10 glucose, 26 NaHCO_3_, 2.5 CaCl_2_ (300 mOsm/L) at 37°C for 15 min. Thereafter, slices were bisected and maintained and recorded at 22°C, with an aCSF flow rate of ∼1.5 ml/min.

For current-clamp recordings and voltage-clamp recordings, the internal pipette solution contained the following (in mM): 130 K-gluconate, 2 KCl, 3 MgCl_2_, 2 MgATP, 0.2 Na_2_GTP, 10 HEPES, 1 EGTA (290 mOsm/L, pH 7.25). To measure spontaneous inhibitory postsynaptic currents (sIPSC) activity, the internal pipette solution contained the following (in mM): 140 KCl, 10 HEPES, 1 EGTA, 5 MgATP, 3.3 MgCl_2_, 0.3 NaGTP (290 mOsm/L, pH 7.25), with 1 mM kynurenic acid added to the aCSF. 0.01% Neurobiotin was included within the pipette solutions for post-processing.

All recordings were acquired with a HEKA EPC10 USB and PATCHMASTER software (HEKA). Voltage-clamp recordings were low pass filtered at 3 kHz and sampled at 7 kHz while current-clamp recordings were filtered at 10 kHz and sampled at 20-25 kHz. We monitored changes in input resistance (R_in_) during the course of the experiments by periodically injecting hyperpolarising steps (−20 pA or −40 pA) in current-clamp mode. Cells were excluded from analysis if R_in_ deviated by >20% over time. For current-clamp recordings, BPN neurons were recorded at their resting membrane potential (RMP). For voltage-clamp recordings, V_h_ was −60 mV. Membrane potential measurements were not corrected for the liquid junction potential between pipette solution and the recording chamber. The reference electrode was a Ag/AgCl^-^ pellet. Only one BPN neuron per slice was recorded during pharmacology experiments. For post-patch processing, slices were fixed overnight in 4% PFA, washed three times in PBS, mounted on SuperFrost slides, and coverslipped with VECTASHIELD mounting medium with DAPI (Vector Laboratories).

### Quantification and Statistical Analysis

In the beam walk task, hindlimb slips were counted manually and traversal time measured by a stopwatch by observers blinded to genotype and treatment. Hindlimb incoordination was likewise evaluated by blinded observers, rated as either present or absent (not graded). For behavioural data and muscle histology, two-way group comparisons were evaluated by two-sided *t*-test; within-subjects comparisons were evaluated by paired *t*-test. Results of the wash-and-bind assay were analyzed by extracting pixel intensity values in each channel for each image (ImageJ). Pixels were thresholded (CB1R intensity >40) and pixel-wise CB1R-COL6A3 intensity correlations were compared between COL6A3^WT^ and COL6A3^ΔC4^ groups by Fisher’s z transformation. For the cAMP assay, the effect of treatment was assessed with ANOVA followed by post-hoc Tukey HSD test for inter-group comparisons. Patch-clamp recording data were analyzed using Clampfit 10 (Molecular Devices) and MiniAnalysis (Synaptosoft). Sample traces were generated with Igor Pro v6 (WaveMetrics) and graphs produced with Prism 7 (GraphPad) or the ggplot2 R package. The nonparametric Kolmogorov–Smirnov test (K–S test; MiniAnalysis) was used to quantify the effects of bath applied pharmacological treatments on each neuron before grouping the data. To compare basal EPSC and IPSC events between genotypes, and the effects of pharmacological interventions, 200 events per neuron per time period in question were used for analysis. To assess effects of pharmacological interventions on voltage-clamp recordings, cells were included for statistical analysis if the peak current was greater than the mean ± 2*SD of the preceding baseline value. Baseline values were taken from a minimum period of 4 min prior to intervention, in 30 s bins. These data were then used to calculate the average frequency, amplitude, and the response compared to baseline (Δ) over time. Unless otherwise stated, data are presented as mean ± SEM with the number of cells per group represented in parentheses for figures. Comparison of pharmacological interventions on the same neuron were analyzed using paired *t*-tests; different groups of cells were compared using unpaired 2-sided *t*-tests. For experiments where data were collected repeatedly for each cell, repeated measures ANOVA was used. A statistical significance threshold of p < 0.05 was used.

## EXTENDED DATA FIGURES

**Figure 2-1.**
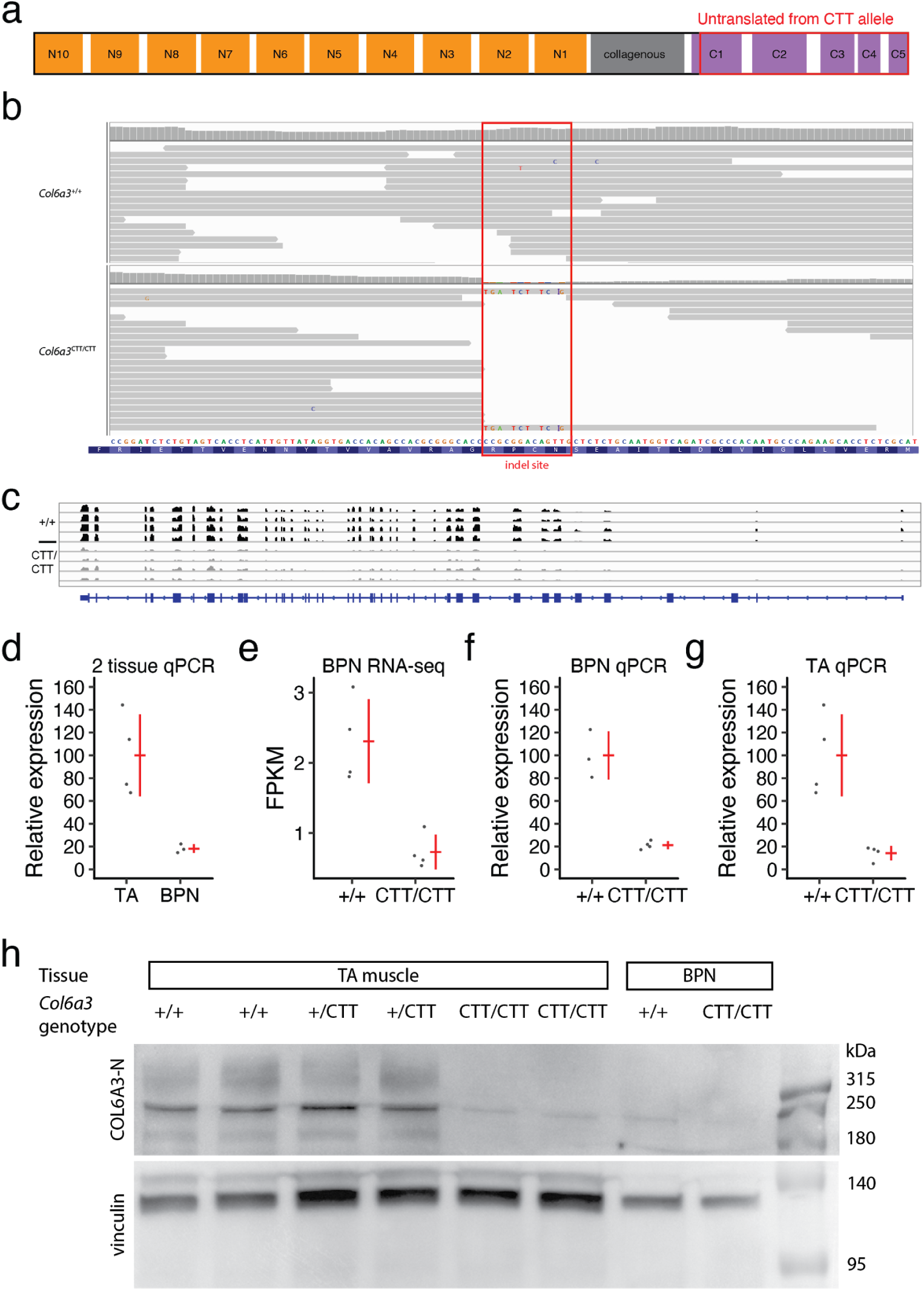
Validation of *Col6a3*^CTT^ allele. **A**, Domains of COL6A3, indicating segment which is untranslated from the *Col6a3*^CTT^ allele. **B**, Aligned BPN RNA-seq reads verify the CTT mutation at the RNA level. **C,** BPN RNA-seq read pileups at the *Col6a3* locus, n=4 replicates per genotype. **D**, Comparison of BPN and tibialis anterior (TA) muscle expression by qPCR. **E-G**, Reduced expression in *Col6a3*^CTT/CTT^ mice in BPN according to RNA-seq (**E**) and qPCR (**F**), and in TA according to qPCR (**G**). **H**, Western blot detection of COL6A3 N-terminal domain in TA muscle and BPN.

**Figure 2-2.**
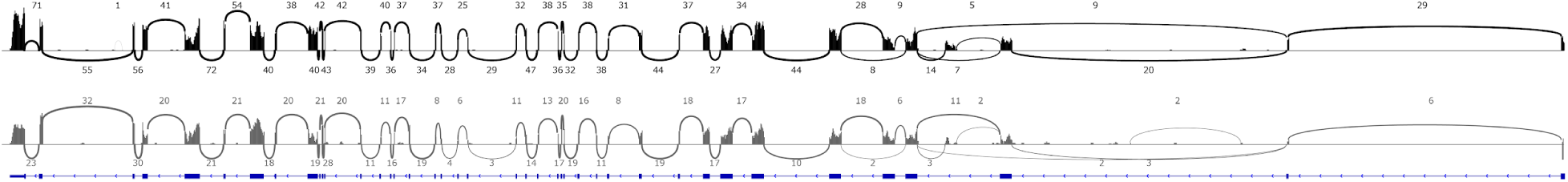
Splicing of *Col6a3* transcripts. Quantification of BPN RNA-seq split reads with 2-pass mapping. Numbers indicate number of reads spanning intron indicated by arc. Top, wildtype; bottom, *Col6a3*^CTT/CTT^. Split reads were consistent with expression of a transcript encoding a full-length COL6A3 protein in wildtypes. In *Col6a3*^CTT/CTT^ mice, exon usage and splicing were indistinguishable from wildtype

**Figure 2-3.**
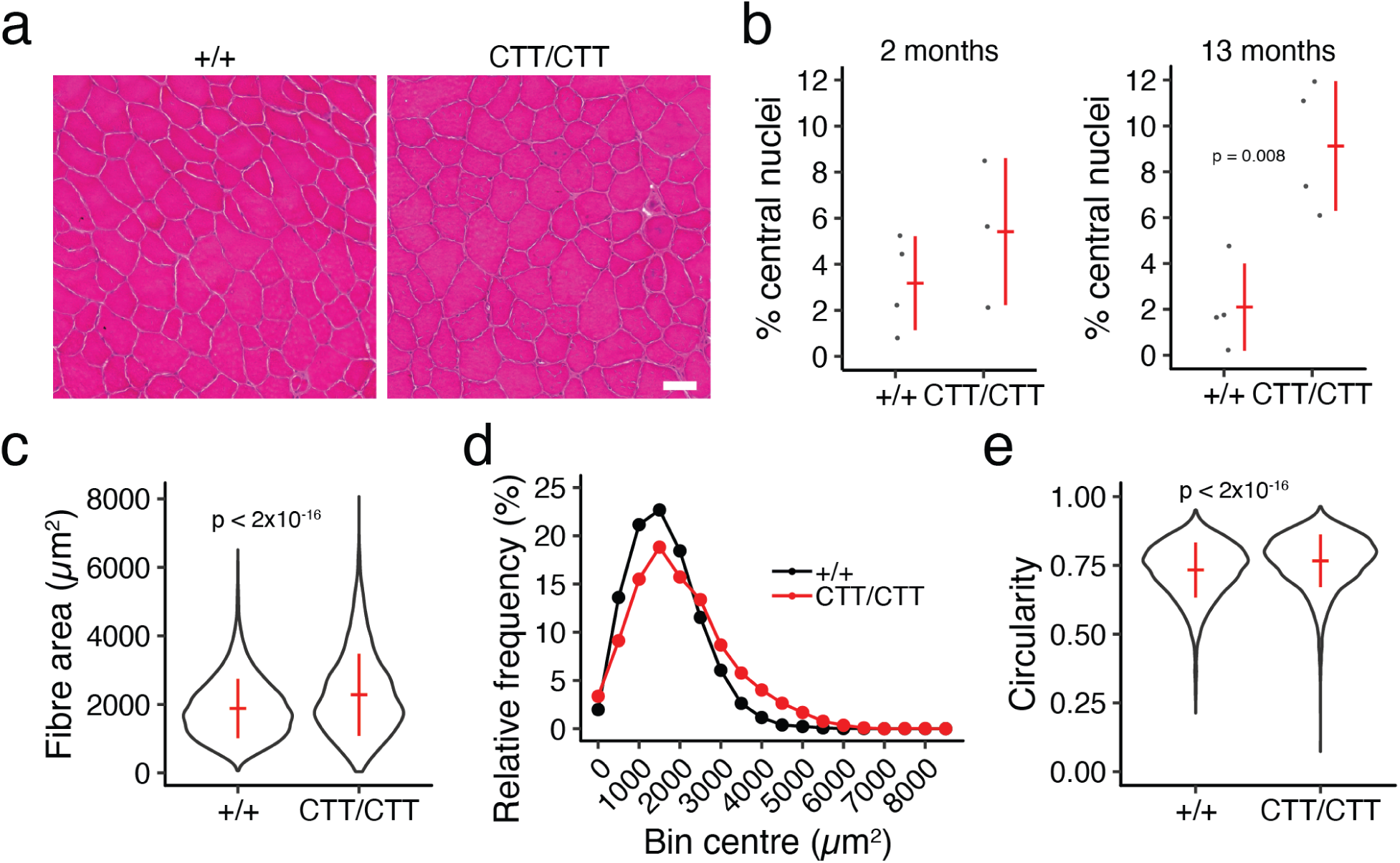
Muscle phenotype of *Col6a3*^CTT/CTT^ mice. **A**, Representative tibialis anterior hematoxylin & eosin staining of tibialis anterior (TA) muscle at 13 months of age. Scale bar, 50 µm. **B**, Proportion of internalised nuclei in TA muscle at age 2 months (n=3-4 per genotype) and 13 months (n=4 per genotype). **C,D**, Quantification (**C**) and distribution (**D**) of TA muscle fibre cross-sectional area in 13 month old mice. **E**, Quantification of circularity of TA muscle fibres in 13 month old mice. n=5,521-5,857 fibres and 4 mice per genotype for c-e. All error bars are standard deviation and centre values are group means. Genotypes were compared by 2-sided *t*-test.

**Figure 2-4.**
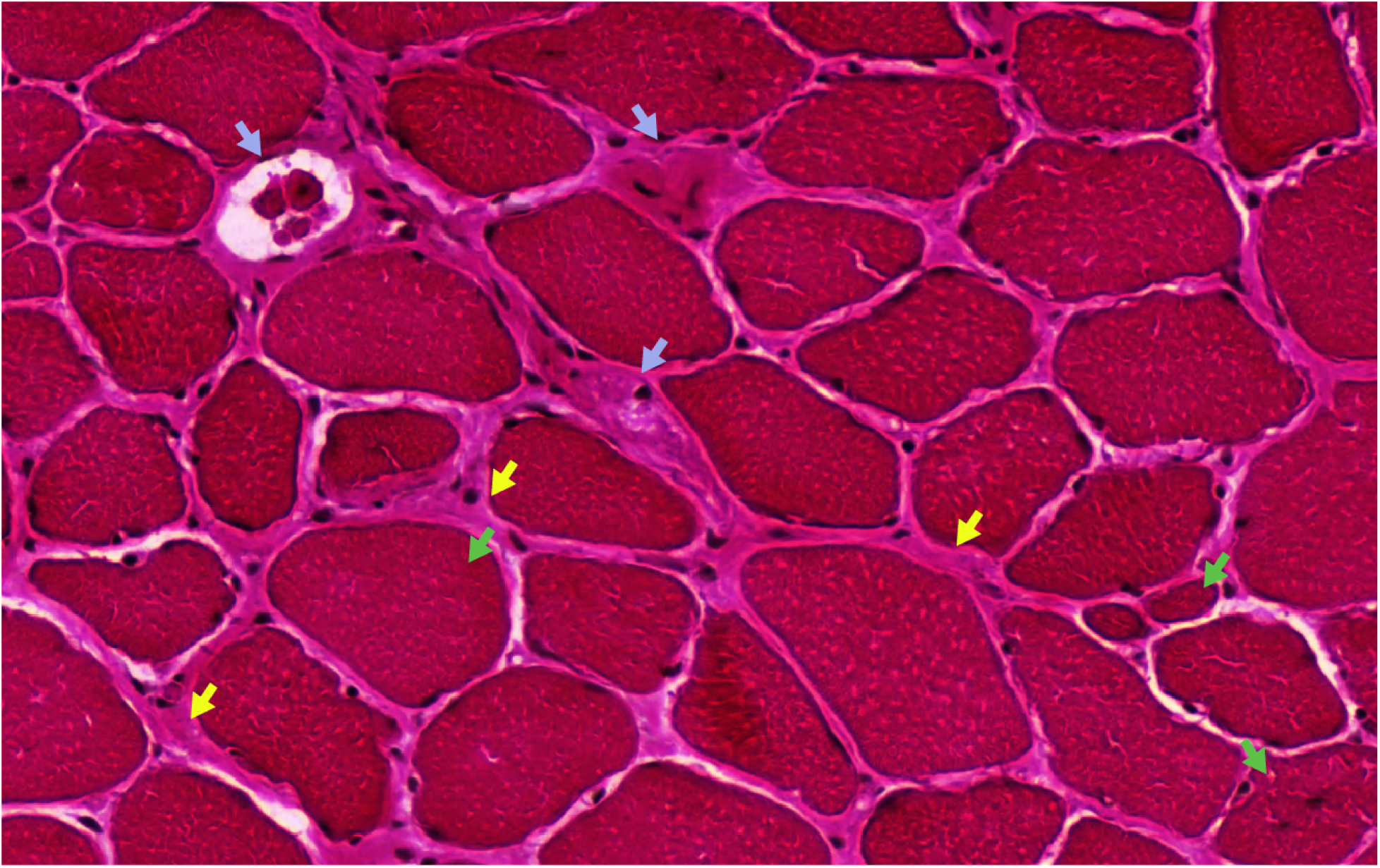
Representative myopathy in aged *Col6a3*^CTT/CTT^ muscle. Hematoxylin & eosin staining of tibialis anterior of 13 month old *Col6a3*^CTT/CTT^ mouse showing endomysial fibrosis (yellow arrows; increased connective tissue between fibres), fibres at different stages of degeneration/necrosis and regeneration (blue arrows), and rounded fibres with increased size variation and/or internalised nuclei (green arrows).

**Figure 3-1.**
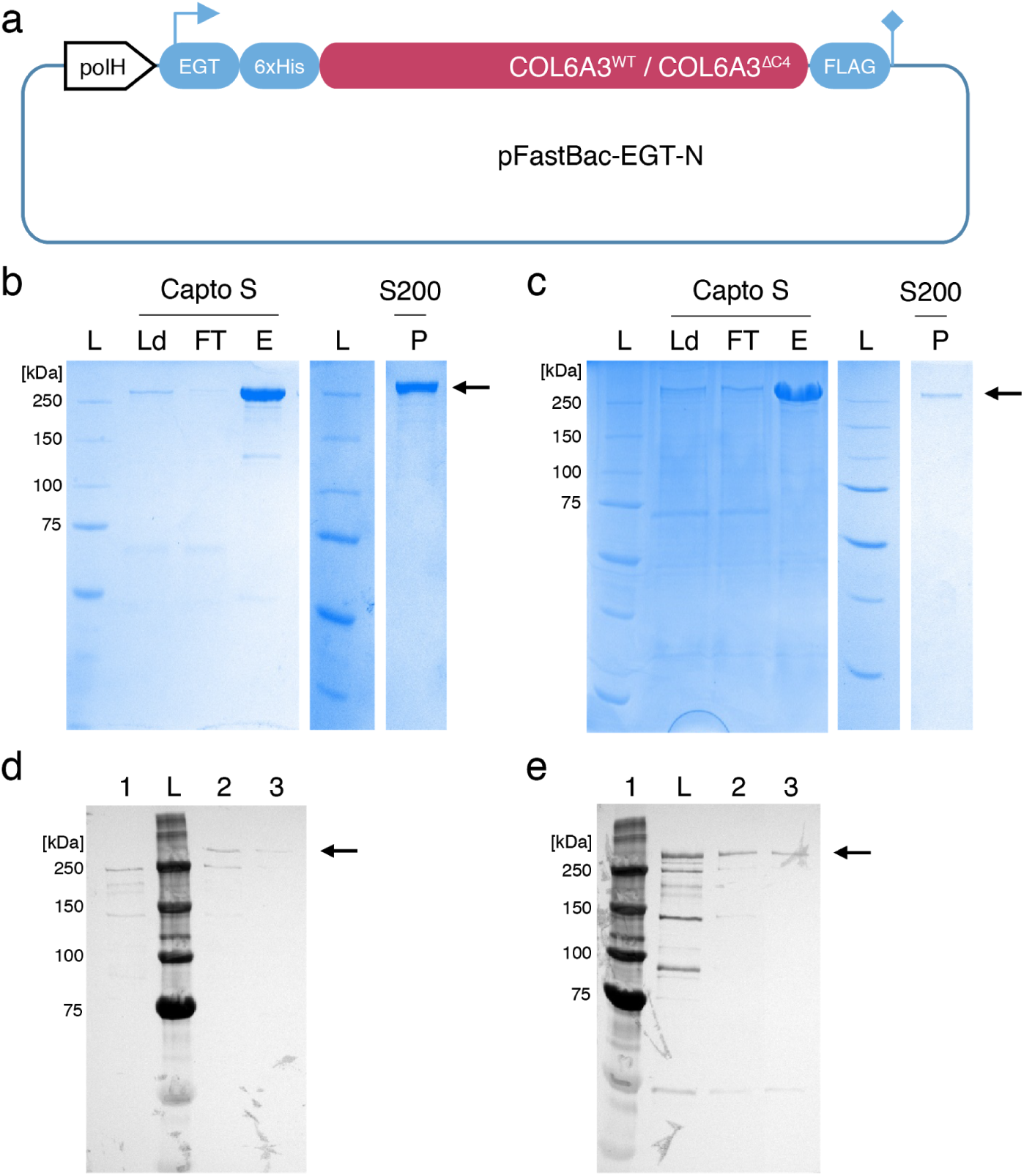
Production of tagged COL6A3 protein. **a**, Schematic overview of the transfer vector pFastBac-EGT-N. COL6A3^WT^ or COL6A3^ΔC4^ expression is driven by the polyhedrine promoter (polH). The main features include the N-terminal secretion signal ecdysteroid UDP-glucosyltransferase (EGT), N-terminal 6xHis tag, and C-terminal FLAG tag. **b,c**, Purification of COL6A3^WT^ (**b**) and COL6A3^ΔC4^ (**c**) by a two step strategy. Collagens were captured from culture medium using cation exchange chromatography (CEX) and polished by size-exclusion chromatography (SEC). 15 µl samples of each purification step were subjected to SDS-PAGE and visualized by Coomassie staining. Arrows indicate the presence of COL6A3 with a predicted weight of 290 kDa (COL6A3^WT^) or 280 kDa (COL6A3^ΔC4^). Capto S, HiTrap Capto S (CEX); S200, S200pg (SEC); L, protein ladder; Ld, load (growth medium); FT, flow-through; E, elution; P, peak (main protein peak from SEC. **d,e**, The success of the two-step purification strategy of COL6A3^WT^ (**d**) and COL6A3^ΔC4^ (**e**) was further validated by Western blotting using a peroxidase-labeled anti-6xHis antibody. 15 µl samples of each purification step were subjected to SDS-PAGE and Western blotting. Arrows indicate the presence of COL6A3 with a predicted weight of 290 kDa (COL6A3^WT^) or 280 kDa (COL6A3^ΔC4^). L, protein ladder; 1, load (growth medium); 2, elution from CEX (HiTrap Capto S); 3, main protein peak from SEC (S200 pg).

**Figure 5-1.**
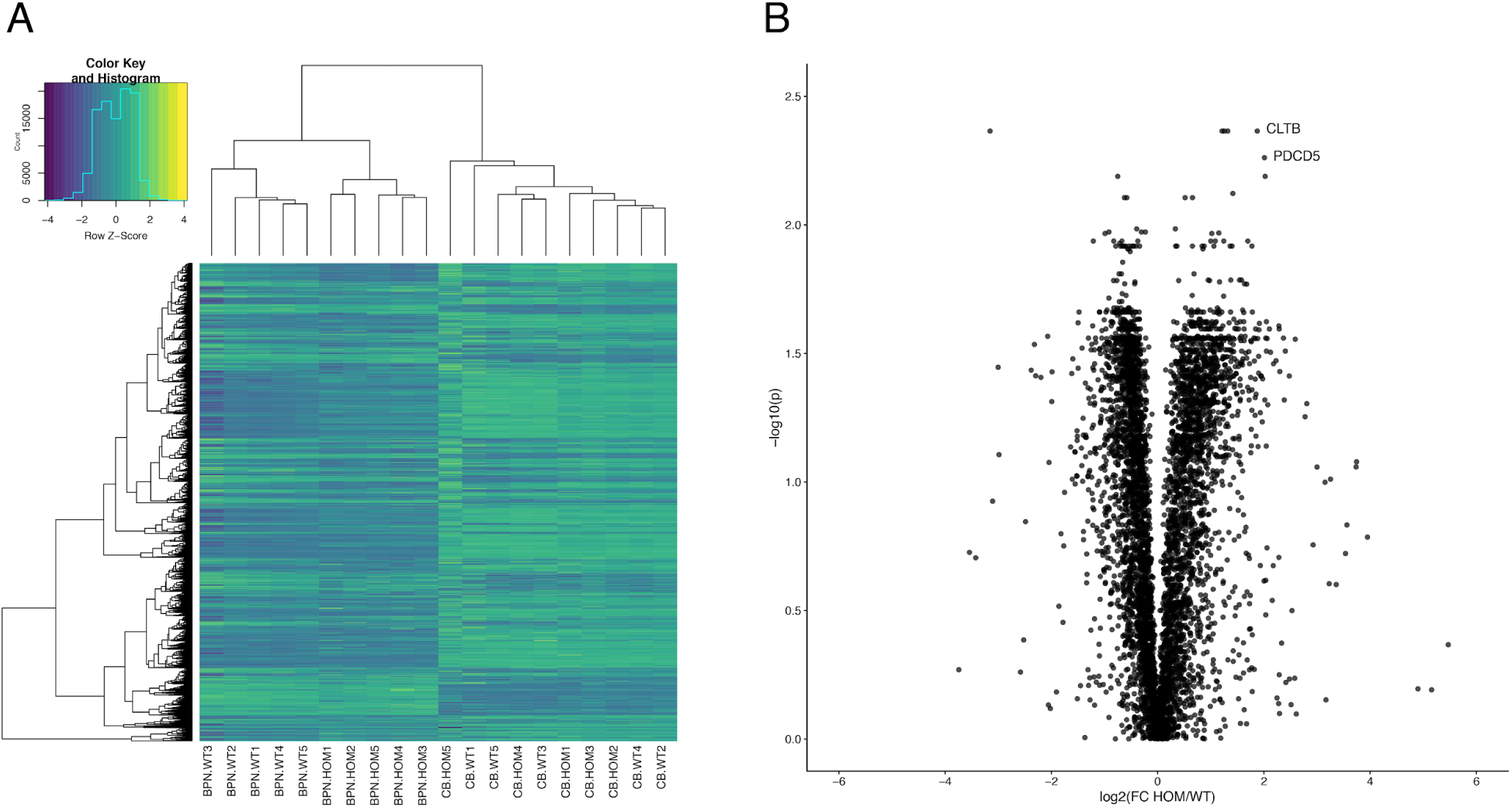
Proteomic analysis of *Col6a3*^CTT/CTT^ brain tissue. **a**, Hierarchical clustering of 5,609 proteins detected my LC-MSMS in BPN and cerebellar (CB) tissue of wildtype (WT) and *Col6a3*^CTT/CTT^ mice (HOM). Genotype clearly separated the samples in the BPN but not CB. **b**, Volcano plot of the BPN proteins, showing genotype differences. CLTB (clathrin B) and PDCD5 (programmed cell death 5) are highlighted. *P* values were adjusted by the false discovery rate method.

**Figure 7-1.**
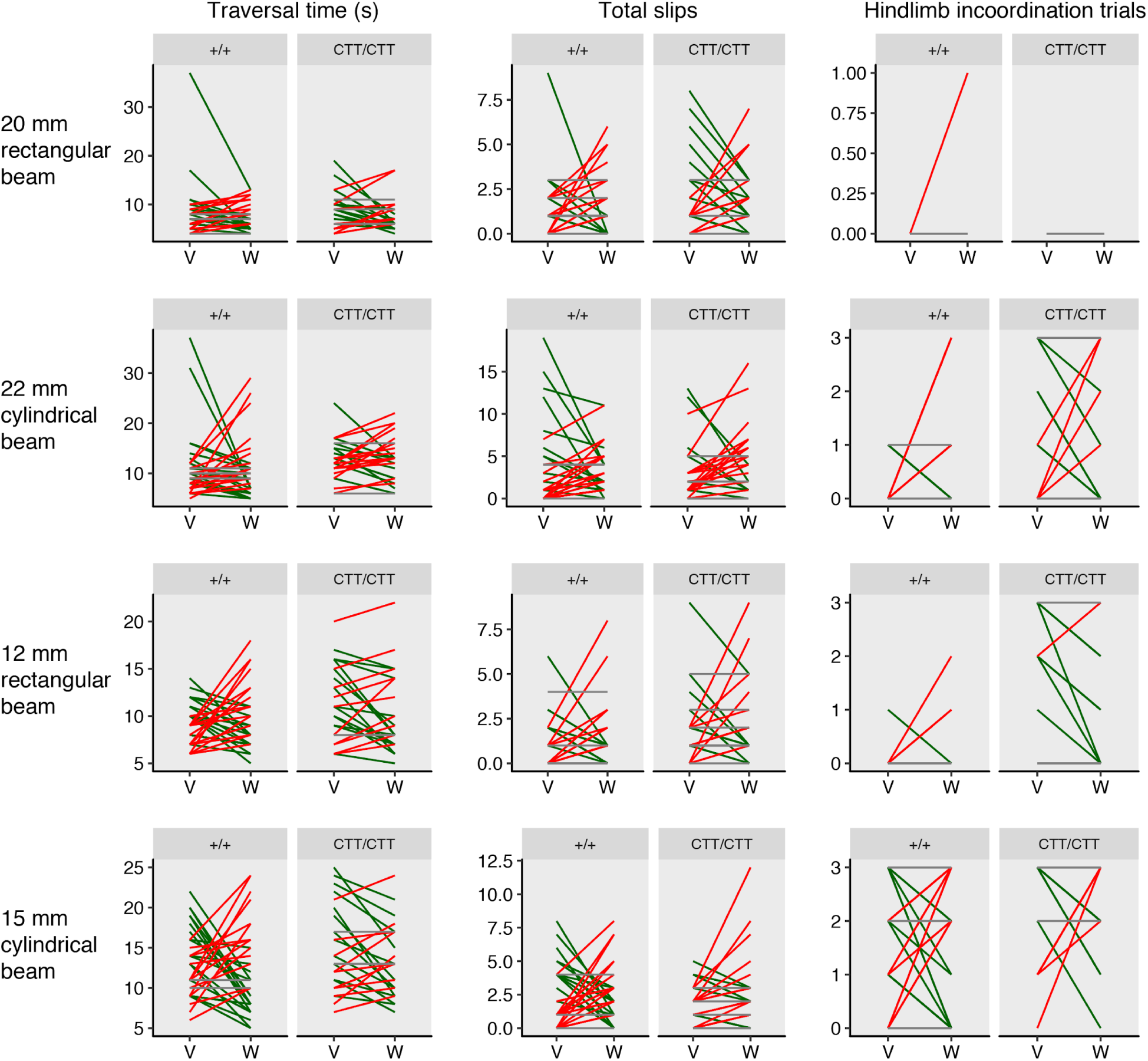
Full results of cannabinoid augmentation experiment. Effect of WIN 55,212-2 (WIN; 1 mg/kg i.p.) treatment on beam walk performance. Each mouse received both WIN and vehicle in a randomized crossover design, with the two conditions separated by 1-8 weeks. Testing commenced 2 hours after injection.Green, performance improved by WIN; red, performance worsened with WIN; grey, performance unchanged with WIN. Male and female mice aged 12-28 wk, n=26-34 per genotype.

## EXTENDED DATA TABLE

**Table 5-1.** Membrane properties and action potential properties (in response to +200 pA ramp; n=9 for all except for AP threshold) of recorded BPN neurons. Values shown are mean ± standard error. RMP, resting membrane potential. AP, action potential.

|  | *Col6a3*<sup>+/+</sup> | *Col6a3*<sup>CTT/CTT</sup> |
| --- | --- | --- |
| **MEMBRANE PROPERTIES** |  |  |
| Capacitance (pF) | 54.74 ± 6.73 (n=19) | 63.64 ± 7.66 (n=19) |
| Resistance (MΩ) | 29.48 ± 3.56 (n=19) | 37.81 ± 4.58 (n=20) |
| RMP (mV) | -66.49 ± 1.16 (n=19) | -66.24 ± 1.60 (n=20) |
| **ACTION POTENTIAL PROPERTIES** |  |  |
| AP threshold (mV) | -50.25 ± 1.59 (n=16) | -48.99 ± 1.45 (n=16) |
| Amplitude (mV) | 78.17 ± 6.50 | 83.32 ± 4.16 |
| Noise (mV) | 0.016 ± 0.005 | 0.009 ± 0.01 |
| Rise (ms) | 7.77 ± 0.21 | 6.71 ± 0.55 |
| Decay (ms) | 0.84 ± 0.10 | 0.81 ± 0.12 |
| Area (mVms) | 111.9 ± 11.46 | 104.7 ± 8.33 |
| Baseline (mV) | -39.56 ± 1.25 | -37.77 ± 2.36 |
| Rise 10-90 (ms) | $1.46 \pm 0.25$ | $1.24 \pm 0.37$ |
| Half-width (ms) | $1.09 \pm 0.11$ | $1.00 \pm 0.13$ |
| Rise 50 (ms) | $0.48 \pm 0.04$ | $0.43 \pm 0.04$ |
| Slope 10-90 (mV/ms) | $61.09 \pm 10.64$ | $87.79 \pm 16.01$ |
| Number of APs | $28.22 \pm 5.62$ | $22.11 \pm 6.95$ |

## SUPPLEMENTARY VIDEO

**Video 2-1. Beam walk performance of *Col6a3*^CTT/CTT^ mice.** Representative beam walk performance showing characteristic hindlimb incoordination in *Col6a3*^CTT/CTT^ mice. Top, wildtype mouse. Bottom, *Col6a3*^CTT/CTT^ mouse with hindlimb incoordination.

## REFERENCES

Albanese, A., Bhatia, K., Bressman, S.B., Delong, M.R., Fahn, S., Fung, V.S.C., Hallett, M., Jankovic, J., Jinnah, H.A., Klein, C., et al. (2013). Phenomenology and classification of dystonia: a consensus update. Mov. Disord. Off. J. Mov. Disord. Soc. 28, 863–873.

Birnboim, H.C., and Doly, J. (1979). A rapid alkaline extraction procedure for screening recombinant plasmid DNA. Nucleic Acids Res. 7, 1513–1523.

Bonaldo, P., and Colombatti, A. (1989). The carboxyl terminus of the chicken alpha 3 chain of collagen VI is a unique mosaic structure with glycoprotein Ib-like, fibronectin type III, and Kunitz modules. J. Biol. Chem. 264, 20235–20239.

Bönnemann, C.G. (2011). The collagen VI-related myopathies: muscle meets its matrix. Nat. Rev. Neurol. 7, 379–390.

Briñas, L., Richard, P., Quijano-Roy, S., Gartioux, C., Ledeuil, C., Lacène, E., Makri, S., Ferreiro, A., Maugenre, S., Topaloglu, H., et al. (2010). Early onset collagen VI myopathies: Genetic and clinical correlations. Ann. Neurol. 68, 511–520.

Brodal, P., and Bjaalie, J.G. (1997). Salient anatomic features of the cortico-ponto-cerebellar pathway. Prog. Brain Res. 114, 227–249.

Brosch, M., Yu, L., Hubbard, T., and Choudhary, J. (2009). Accurate and sensitive peptide identification with Mascot Percolator. J. Proteome Res. 8, 3176–3181.

Bruderer, R., Bernhardt, O.M., Gandhi, T., Miladinović, S.M., Cheng, L.-Y., Messner, S., Ehrenberger, T., Zanotelli, V., Butscheid, Y., Escher, C., et al. (2015). Extending the limits of quantitative proteome profiling with data-independent acquisition and application to acetaminophen-treated three-dimensional liver microtissues. Mol. Cell. Proteomics MCP 14, 1400–1410.

Calabresi, P., Pisani, A., Rothwell, J., Ghiglieri, V., Obeso, J.A., and Picconi, B. (2016). Hyperkinetic disorders and loss of synaptic downscaling. Nat. Neurosci. 19, 868–875.

Cescon, M., Gattazzo, F., Chen, P., and Bonaldo, P. (2015). Collagen VI at a glance. J. Cell Sci. 128, 3525–3531.

Cescon, M., Chen, P., Castagnaro, S., Gregorio, I., and Bonaldo, P. (2016). Lack of collagen VI promotes neurodegeneration by impairing autophagy and inducing apoptosis during aging. Aging 8, 1083–1098.

Cheng, J.S., Dubal, D.B., Kim, D.H., Legleiter, J., Cheng, I.H., Yu, G.-Q., Tesseur, I., Wyss-Coray, T., Bonaldo, P., and Mucke, L. (2009). Collagen VI protects neurons against Aβ toxicity. Nat. Neurosci. 12, 119.

Chevaleyre, V., and Castillo, P.E. (2003). Heterosynaptic LTD of hippocampal GABAergic synapses: a novel role of endocannabinoids in regulating excitability. Neuron 38, 461–472.

Chopra, R.K., and Ananthanarayanan, V.S. (1982). Conformational implications of enzymatic proline hydroxylation in collagen. Proc. Natl. Acad. Sci. U. S. A. 79, 7180–7184.

Concordet, J.-P., and Haeussler, M. (2018). CRISPOR: intuitive guide selection for CRISPR/Cas9 genome editing experiments and screens. Nucleic Acids Res. 46, W242–W245.

Demir, E., Sabatelli, P., Allamand, V., Ferreiro, A., Moghadaszadeh, B., Makrelouf, M., Topaloglu, H., Echenne, B., Merlini, L., and Guicheney, P. (2002). Mutations in COL6A3 cause severe and mild phenotypes of Ullrich congenital muscular dystrophy. Am. J. Hum. Genet. 70, 1446–1458.

Dobin, A., Davis, C.A., Schlesinger, F., Drenkow, J., Zaleski, C., Jha, S., Batut, P., Chaisson, M., and Gingeras, T.R. (2013). STAR: ultrafast universal RNA-seq aligner. Bioinformatics 29, 15–21.

Folker, E.S., and Baylies, M.K. (2013). Nuclear positioning in muscle development and disease. Front. Physiol. 4.

Frik, J., Merl-Pham, J., Plesnila, N., Mattugini, N., Kjell, J., Kraska, J., Gómez, R.M., Hauck, S.M., Sirko, S., and Götz, M. (2018). Cross-talk between monocyte invasion and astrocyte proliferation regulates scarring in brain injury. EMBO Rep. 19.

Grosche, A., Hauser, A., Lepper, M.F., Mayo, R., von Toerne, C., Merl-Pham, J., and Hauck, S.M. (2016). The Proteome of Native Adult Müller Glial Cells From Murine Retina. Mol. Cell. Proteomics MCP 15, 462–480.

Grumati, P., Coletto, L., Sabatelli, P., Cescon, M., Angelin, A., Bertaggia, E., Blaauw, B., Urciuolo, A., Tiepolo, T., Merlini, L., et al. (2010). Autophagy is defective in collagen VI muscular dystrophies, and its reactivation rescues myofiber degeneration. Nat. Med. 16, 1313–1320.

Guo, J.-Z., Sauerbrei, B., Cohen, J.D., Mischiati, M., Graves, A., Pisanello, F., Branson, K., and Hantman, A.W. (2019). The Pontine Nuclei are an Integrative Cortico-Cerebellar Link Critical for Dexterity. BioRxiv 637447.

Hauck, S.M., Dietter, J., Kramer, R.L., Hofmaier, F., Zipplies, J.K., Amann, B., Feuchtinger, A., Deeg, C.A., and Ueffing, M. (2010). Deciphering membrane-associated molecular processes in target tissue of autoimmune uveitis by label-free quantitative mass spectrometry. Mol. Cell. Proteomics MCP 9, 2292–2305.

Heumüller, S.E., Talantikite, M., Napoli, M., Armengaud, J., Mörgelin, M., Hartmann, U., Sengle, G., Paulsson, M., Moali, C., and Wagener, R. (2019). C-terminal proteolysis of the collagen VI α3 chain by BMP-1 and proprotein convertase(s) releases endotrophin in fragments of different sizes. J. Biol. Chem. 294, 13769–13780.

Hua, T., Vemuri, K., Nikas, S.P., Laprairie, R.B., Wu, Y., Qu, L., Pu, M., Korde, A., Jiang, S., Ho, J.-H., et al. (2017). Crystal structures of agonist-bound human cannabinoid receptor CB1. Nature 547, 468–471.

Irwin, W.A., Bergamin, N., Sabatelli, P., Reggiani, C., Megighian, A., Merlini, L., Braghetta, P., Columbaro, M., Volpin, D., Bressan, G.M., et al. (2003). Mitochondrial dysfunction and apoptosis in myopathic mice with collagen VI deficiency. Nat. Genet. 35, 367–371.

Kelly, R.M., and Strick, P.L. (2003). Cerebellar loops with motor cortex and prefrontal cortex of a nonhuman primate. J. Neurosci. Off. J. Soc. Neurosci. 23, 8432–8444.

Koppel, B.S. (2015). Cannabis in the Treatment of Dystonia, Dyskinesias, and Tics. Neurother. J. Am. Soc. Exp. Neurother. 12, 788–792.

Kreitzer, A.C., and Malenka, R.C. (2007). Endocannabinoid-mediated rescue of striatal LTD and motor deficits in Parkinson’s disease models. Nature 445, 643–647.

Kushmerick, C., Price, G.D., Taschenberger, H., Puente, N., Renden, R., Wadiche, J.I., Duvoisin, R.M., Grandes, P., and von Gersdorff, H. (2004). Retroinhibition of presynaptic Ca2+ currents by endocannabinoids released via postsynaptic mGluR activation at a calyx synapse. J. Neurosci. Off. J. Soc. Neurosci. 24, 5955–5965.

Lepper, M.F., Ohmayer, U., von Toerne, C., Maison, N., Ziegler, A.-G., and Hauck, S.M. (2018). Proteomic Landscape of Patient-Derived CD4+ T Cells in Recent-Onset Type 1 Diabetes. J. Proteome Res. 17, 618–634.

Lo, H.-R., and Chao, Y.-C. (2004). Rapid titer determination of baculovirus by quantitative real-time polymerase chain reaction. Biotechnol. Prog. 20, 354–360.

Marakhonov, A.V., Tabakov, V.Y., Zernov, N.V., Dadali, E.L., Sharkova, I.V., and Skoblov, M.Y. (2018). Two novel COL6A3 mutations disrupt extracellular matrix formation and lead to myopathy from Ullrich congenital muscular dystrophy and Bethlem myopathy spectrum. Gene 672, 165–171.

Marsicano, G., Goodenough, S., Monory, K., Hermann, H., Eder, M., Cannich, A., Azad, S.C., Cascio, M.G., Gutiérrez, S.O., van der Stelt, M., et al. (2003). CB1 cannabinoid receptors and on-demand defense against excitotoxicity. Science 302, 84–88.

Mattugini, N., Merl-Pham, J., Petrozziello, E., Schindler, L., Bernhagen, J., Hauck, S.M., and Götz, M. (2018). Influence of white matter injury on gray matter reactive gliosis upon stab wound in the adult murine cerebral cortex. Glia 66, 1644–1662.

Morgese, M.G., Cassano, T., Cuomo, V., and Giuffrida, A. (2007). Anti-dyskinetic effects of cannabinoids in a rat model of Parkinson’s disease: role of CB(1) and TRPV1 receptors. Exp. Neurol. 208, 110–119.

Pan, T.-C., Zhang, R.-Z., Markova, D., Arita, M., Zhang, Y., Bogdanovich, S., Khurana, T.S., Bönnemann, C.G., Birk, D.E., and Chu, M.-L. (2013). COL6A3 protein deficiency in mice leads to muscle and tendon defects similar to human collagen VI congenital muscular dystrophy. J. Biol. Chem. 288, 14320–14331.

Pan, T.-C., Zhang, R.-Z., Arita, M., Bogdanovich, S., Adams, S.M., Gara, S.K., Wagener, R., Khurana, T.S., Birk, D.E., and Chu, M.-L. (2014). A mouse model for dominant collagen VI disorders: heterozygous deletion of Col6a3 Exon 16. J. Biol. Chem. 289, 10293–10307.

Pappas, S.S., Li, J., LeWitt, T.M., Kim, J.-K., Monani, U.R., and Dauer, W.T. (2018). A cell autonomous torsinA requirement for cholinergic neuron survival and motor control. ELife 7.

Quartarone, A., and Pisani, A. (2011). Abnormal plasticity in dystonia: Disruption of synaptic homeostasis. Neurobiol. Dis. 42, 162–170.

Robbe, D., Kopf, M., Remaury, A., Bockaert, J., and Manzoni, O.J. (2002). Endogenous cannabinoids mediate long-term synaptic depression in the nucleus accumbens. Proc. Natl. Acad. Sci. U. S. A. 99, 8384–8388.

Schindelin, J., Arganda-Carreras, I., Frise, E., Kaynig, V., Longair, M., Pietzsch, T., Preibisch, S., Rueden, C., Saalfeld, S., Schmid, B., et al. (2012). Fiji: an open-source platform for biological-image analysis. Nat. Methods 9, 676–682.

Shimomura, H., Lee, T., Tanaka, Y., Awano, H., Itoh, K., Nishino, I., and Takeshima, Y. (2019). Two closely spaced mutations in cis result in Ullrich congenital muscular dystrophy. Hum. Genome Var. 6, 1–4.

Suzuki, L., Coulon, P., Sabel-Goedknegt, E.H., and Ruigrok, T.J.H. (2012). Organization of cerebral projections to identified cerebellar zones in the posterior cerebellum of the rat. J. Neurosci. Off. J. Soc. Neurosci. 32, 10854–10869.

Timpl, R., and Chu, M. (1994). Microfibrillar collagen type VI. In Extracellular Matrix Assembly and Structure, (Academic Press), pp. 207–242.

UniProt Consortium (2019). UniProt: a worldwide hub of protein knowledge. Nucleic Acids Res. 47, D506–D515.

Urciuolo, A., Quarta, M., Morbidoni, V., Gattazzo, F., Molon, S., Grumati, P., Montemurro, F., Tedesco, F.S., Blaauw, B., Cossu, G., et al. (2013). Collagen VI regulates satellite cell self-renewal and muscle regeneration. Nat. Commun. 4, 1964.

Varma, N., Carlson, G.C., Ledent, C., and Alger, B.E. (2001). Metabotropic glutamate receptors drive the endocannabinoid system in hippocampus. J. Neurosci. Off. J. Soc. Neurosci. 21, RC188.

Vitagliano, L., Berisio, R., Mazzarella, L., and Zagari, A. (2001). Structural bases of collagen stabilization induced by proline hydroxylation. Biopolymers 58, 459–464.

Wagner, M.J., Kim, T.H., Kadmon, J., Nguyen, N.D., Ganguli, S., Schnitzer, M.J., and Luo, L. (2019). Shared Cortex-Cerebellum Dynamics in the Execution and Learning of a Motor Task. Cell.

Wilson, R.I., and Nicoll, R.A. (2001). Endogenous cannabinoids mediate retrograde signalling at hippocampal synapses. Nature 410, 588–592.

Wiśniewski, J.R., Zougman, A., Nagaraj, N., and Mann, M. (2009). Universal sample preparation method for proteome analysis. Nat. Methods 6, 359–362.

Xiao, T., and Baier, H. (2007). Lamina-specific axonal projections in the zebrafish tectum require the type IV collagen Dragnet. Nat. Neurosci. 10, 1529–1537.

Xiao, T., Staub, W., Robles, E., Gosse, N.J., Cole, G.J., and Baier, H. (2011). Assembly of lamina-specific neuronal connections by slit bound to type IV collagen. Cell 146, 164–176.

Zech, M., Lam, D.D., Francescatto, L., Schormair, B., Salminen, A.V., Jochim, A., Wieland, T., Lichtner, P., Peters, A., Gieger, C., et al. (2015). Recessive mutations in the α3 (VI) collagen gene COL6A3 cause early-onset isolated dystonia. Am. J. Hum. Genet. 96, 883–893.

Zeisel, A., Hochgerner, H., Lönnerberg, P., Johnsson, A., Memic, F., van der Zwan, J., Häring, M., Braun, E., Borm, L.E., La Manno, G., et al. (2018). Molecular Architecture of the Mouse Nervous System. Cell 174, 999–1014.e22.

Zhang, Y.-Z., Zhao, D.-H., Yang, H.-P., Liu, A.-J., Chang, X.-Z., Hong, D.-J., Bonnemann, C., Yuan, Y., Wu, X.-R., and Xiong, H. (2014). Novel collagen VI mutations identified in Chinese patients with Ullrich congenital muscular dystrophy. World J. Pediatr. WJP 10, 126–132.

